# Glycoproteogenomics characterizes the CD44 splicing code driving bladder cancer invasion

**DOI:** 10.1101/2021.09.04.458979

**Authors:** Cristiana Gaiteiro, Janine Soares, Marta Relvas-Santos, Andreia Peixoto, Dylan Ferreira, Andreia Brandão, Elisabete Fernandes, Rita Azevedo, Paula Paulo, Carlos Palmeira, Luís Lima, Rui Freitas, Andreia Miranda, Hugo Osório, André M. N. Silva, Jesús Prieto, Lúcio Lara Santos, José Alexandre Ferreira

## Abstract

Bladder cancer (BC) management demands the introduction of novel molecular targets for precision medicine. Cell surface glycoprotein CD44 has been widely studied as a potential biomarker of BC aggressiveness and cancer stem cells. However, significant alternative splicing and multiple glycosylation generate a myriad of glycoproteoforms with potentially distinct functional roles. The lack of tools for precise molecular characterization has led to conflicting results, delaying clinical applications. Addressing these limitations, we have interrogated the transcriptome of a large BC patient cohort for splicing signatures. Remarkable CD44 heterogeneity was observed, as well as associations between short CD44 standard splicing isoform (CD44s), invasion and poor prognosis. In parallel, immunoassays showed that targeting short *O*-glycoforms could hold the key to improve CD44 cancer specificity. This prompted the development of a glycoproteogenomics approach, building on the integration of transcriptomics-customized datasets and glycomics for protein annotation from nanoLC-ESI-MS/MS experiments. The concept was applied to invasive human BC cell lines, glycoengineered cells, and tumor tissues, enabling unequivocal CD44s identification. Finally, we confirmed the link between CD44s and invasion *in vitro* by siRNA knockdown, supporting findings from BC tissues. The key role played by short-chain *O*-glycans in CD44-mediated invasion was also demonstrated through glycoengineered cell models. Overall, CD44s emerged as biomarker of poor prognosis and CD44-Tn/STn as promising molecular signatures for targeted interventions. This study materializes the concept of glycoproteogenomics and provides a key vision to address the cancer splicing code at the protein level, which may now be expanded to better understand CD44 functional role in health and disease.

**Significance Statement:** The biological role of CD44, a cell membrane glycoprotein involved in most cancer hallmarks and widely explored in BC, is intimately linked to its protein isoforms. mRNA alternative splicing generates several closely related polypeptide sequences, which have so far been inferred from transcripts analysis, in the absence of workflows for unequivocal protein annotation. Dense *O*-glycosylation is also key for protein function and may exponentiate the number of proteoforms, rendering CD44 molecular characterization a daunting enterprise. Here, we integrated multiple molecular information (RNA, proteins, glycans) for definitive CD44 characterization by mass spectrometry, materializing the concept of glycoproteogenomics. BC specific glycoproteoforms linked to invasion have been identified, holding potential for precise cancer targeting. The approach may be transferable to other tumors, paving the way for precision oncology.

## Introduction

Bladder cancer (BC) remains a pressing health concern and encompasses significant mortality, especially when diagnosed at advanced stages (1). Cluster of Differentiation 44 (CD44) is a multifunctional and heavily glycosylated transmembrane protein (2) involved in cell-cell and cell-extracellular matrix adhesion (3), immune functions (4, 5), lymphocyte homing (6, 7), hematopoiesis (8), and oncogenic signaling. By interacting with several downstream effector proteins, it dictates cell migration and adhesion (9, 10), tumor invasion (11), and metastasis (12, 13). Hence, this glycoprotein has been found overexpressed in more aggressive bladder tumors (14, 15), being widely adopted as a biomarker of bladder cancer stem cells (CSC) (16, 17).

The human CD44 gene is located on chromosome 11p13 and consists of 19 exons (2, 18); however, the full-length protein has never been detected due to the occurrence of intense alternative splicing. Exons 1 to 16 encode the extracellular domain, exon 17 the transmembrane region, and exons 18 and 19 the intracellular domains (2). The first 5 exons are constitutively transcribed across all currently known isoforms, encoding hyaluronic acid (19), osteopontin (20), collagen (21), laminin (22), and fibronectin (23) binding sites. On the other hand, exons 6 to 14 are subjected to alternative splicing, generating a myriad of different variants whose functional implications are yet to be fully understood (2). The number of proteoforms generated by mRNA translation and processing is greatly amplified by *O*-GalNAc glycosylation, mostly occurring in CD44 variable regions that are rich in serine and threonine residues. Moreover, a multiplicity of different but closely related glycan structures may be found in the same protein, exponentiating molecular micro-, macro-, and meta-heterogeneity (2, 24). Still, few studies have addressed the CD44 glycocode in cancer and its functional implications for disease progression (25-27). Moreover, conflicting results have been generated concerning its role in disease and clinical value, which are directly linked to analytical limitations for unequivocal molecular characterization, mostly due to the lack of high-throughput approaches to characterize CD44 at the protein level (2). Misguiding nomenclature has also been posing a major limitation for comprehensive data mining and definitive molecular characterization, leading us to propose an uniformization (summarized in **SI Appendix, Fig. S1**) (2).

High-throughput proteomics constitutes the gold standard approach to tackle CD44 molecular heterogeneity; however, the success of current workflows is limited by the capacity of databases for protein annotation. The high degree of sequence similarity amongst proteoforms, many times differing by short and potentially heavily glycosylated peptide sequences, poses a significant limitation. As such, CD44 molecular characterization has been mainly inferred from transcripts analysis, supported by immunoassays based on antibodies that lack isoform specificity. In summary, a single omics cannot portrait its molecular complexity, delaying clinical applications. Herein, we hypothesize that multi-omics settings may be required to unequivocally characterize CD44 glycoproteoforms, combining transcriptomics, glycomics and glycoproteomics in glycoproteogenomics settings (28) for precise identification of CD44 signatures of clinical relevance, foreseeing the design of novel targeted therapies.

## Results

The CD44 glycoproteocode remains poorly characterized in BC and other tumors, frustrating expectations for precise clinical interventions. This is closely related to CD44 high molecular heterogeneity, resulting from intense alternative splicing, close sequence similarity between multiple proteoforms, and dense and diverse *O*-glycosylation, amplified by conflicting nomenclature. Herein, we show that multi-omics, combining transcriptomics, glycomics, and glycoproteomics in glycoproteogenomics settings, are required for precise identification of CD44 signatures of clinical relevance.

### High CD44s/st mRNA is associated with muscle invasion and worst prognosis in bladder cancer

We have started by screening the healthy urothelium (11 cases) and a broad series of bladder tumors (34 non-muscle invasive and 41 muscle-invasive bladder tumors) representative of all disease stages for total CD44 mRNA and 5 experimentally confirmed splicing isoforms (*CD44v2-10, CD44v3-10, CD44v8-10, CD44s/st* and *CD44sol)*. For simplification and easy follow-up, we have adopted the nomenclature proposed by Azevedo, R. *et al*. (2), which reflects the nature of the transcribed exons. Tailored-made primers were designed to specifically detect mRNA encoding for the 4 splicing variants (**SI Appendix, Table S1**) which have been found frequently in other cancer models (29-33). Nevertheless, due to high similarity, this approach was unable to differentiate *CD44*s from the close related *CD44st* isoform containing a shorter cytoplasmic tail (**SI Appendix, Fig. S1**).

In general, *CD44* mRNA was expressed more abundantly in the healthy urothelium than in bladder tumors and showed a trend to decrease with the severity of the disease (**Fig. 1A**). Furthermore, healthy tissues and tumors presented mRNAs encoding for all targeted isoforms, supporting high microheterogeneity. Still, *CD44sol* mRNA was either vestigial or undetected in most samples and, therefore, has not been represented in **Fig. 1A**. Zooming in on the nature of the other transcripts, we found that superficial tumors were enriched for longer isoforms (*CD44v2-10 and/or CD44v3-10*) in comparison to muscle-invasive tumors (**Fig. 1A**). Moreover, invasive tumors expressed higher percentages of shorter CD44 mRNAs lacking the variable region (*CD44s/st*). We then comprehensively interrogated a larger and more homogeneous patient series comprehending over 400 muscle-invasive tumors from The Cancer Genome Atlas (TCGA) with a detailed clinical history (**Figs. 1B-D**). Whole transcriptome analysis confirmed the presence of previously studied isoforms plus *CD44v10* in muscle-invasive tumors. *In silico* analysis revealed that late-stage invasive disease (>T2) and more aggressive non-papillary lesions (34), presented significantly higher levels of short *CD44* transcripts (*CD44sol; CD44s; CD44v10*) in comparison to T2 (**Fig. 1B**) and papillary tumors (**SI Appendix, Fig. S2**), respectively. Furthermore, high levels of *CD44*s (*p*=0.029) and *CD44*v10 (*p*=0.049) were significantly associated with decreased overall survival (**Fig. 1C**). However, none of them were independent predictors of poor prognosis when adjusted to disease stage, which was a relevant variable in this context.

**Fig. 1.**
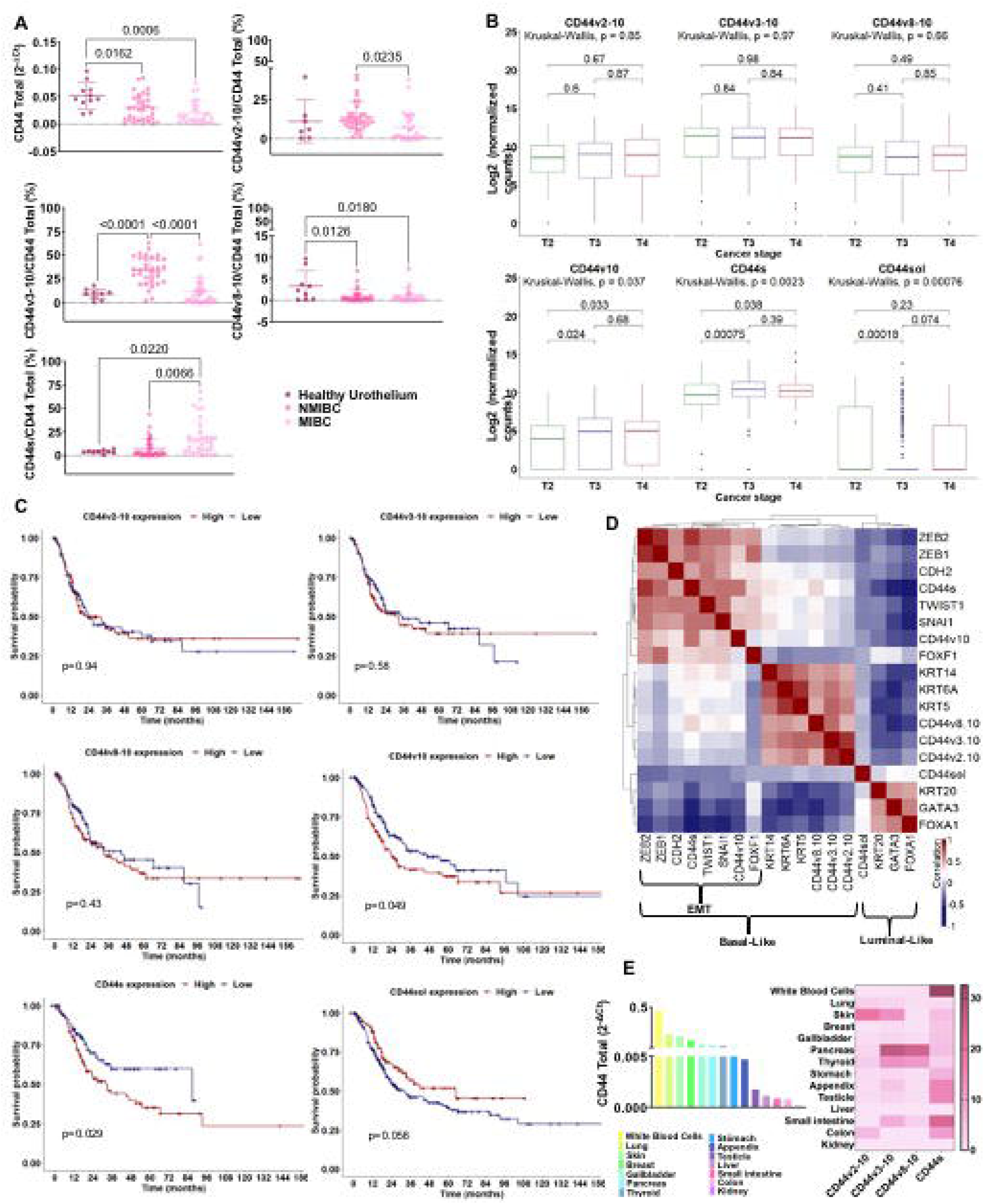
*CD44s* is overexpressed in more aggressive bladder tumors, is not present in the healthy urothelium and shows limited expression in other healthy human tumors. **A) Expression of CD44 and its variants in the healthy urothelium and bladder tumors**. *CD44* gene expression significantly decreased in bladder tumors in comparison to the healthy urothelium. Furthermore, the nature of the variants changed with the severity of disease, with non-muscle invasive bladder cancer (NMIBC), showing higher expression of lengthier isoforms (*CD44v2-10*; *CD44v3-10*) and a significantly lower abundance of CD44s in comparison to muscle-invasive bladder cancer (MIBC). MIBC presented an opposite pattern, supporting enrichment for shorter CD44s isoform. The *CD44sol* isoform was also evaluated but its expression was vestigial and not linked to any tumor type, as such it was not represented in the panel A. **B) TCGA analysis of 413 MIBC cases confirmed the association of CD44s with more aggressive late-stage disease**. Briefly, *CD44s* mRNA was elevated in T3/4 tumors in comparison to T2 tumors, supporting its association with invasion, confirming the observations from our patient’s dataset. **C) Elevated *CD44v10* and *CD44s* mRNA significantly associate with worst prognosis in MIBC**. The Kaplan-Meier curves highlight a clear link between the expression of shorter CD44 isoforms and worst prognosis in bladder cancer. Also, despite its low expression, patients with tumors presenting high *CD44sol* mRNA presented better prognosis. **D) CD44s is expressed by a subgroup of basal-like tumors enriched for genes defining mesenchymal traits**. In general, *CD44sol* associated with more differentiated and less aggressive luminal tumors, whereas the other forms of CD44 related with basal phenotypes, frequently less cohesive and poorly differentiated lesions. However, *CD44s and CD44v10* were characteristic of a subgroup of basal tumors enriched for *ZEB1/2, CDH2, TWIST1, SNAI1* that define mesenchymal phenotypes. Collectively, these observations link shorter CD44 isoforms to bladder cancer invasion and poor prognosis. **E) Total *CD44* and isoforms expressions in relevant healthy cells and organs**. The *CD44* gene presents a heterogeneous expression pattern in healthy tissues, and multiple isoforms may coexist in the same organ. Notably, *CD44s* was expressed by most of the studied tissues, showing significantly high mRNA levels in white blood cells. The gastrointestinal and colorectal tracts as well as the testicle also present high *CD44s*, however, with low total *CD44* levels. Collectively, this result shows that CD44s is not a cancer specific signature. The results correspond to the mean and standard deviation for three independent experiments. Triplicate measurements were conducted for each experiment. P values are presented for one-way ANOVA, nonparametric Wilcoxon, and Kruskal-Wallis tests.

Finally, CD44 transcripts were evaluated in the context of transcriptome-based molecular subtypes of BC. Namely, the luminal subtype, with favorable prognosis, overexpressing *KRT20* as well as *FOXA1*, and *GATA3*, and the basal-like subtype, displaying worst prognosis, high *KRT5* and/or *KRT14* and/or *KRT6A* (35, 36). In the analysis, we also included known epithelial-to-mesenchymal transition (EMT) markers closely linked to invasion (*FOXF1, CDH2, ZEB1, ZEB2, SNAI1, TWIST1*). The Spearman correlation plot in **Fig. 1D** highlights three clear clusters corresponding to these groups. Some proximity between the basal-like and EMT molecular phenotypes could also be observed, supporting the existence of a basal subgroup enriched for more invasive traits. The luminal molecular subtype was not characterized by *CD44* expression. On the other hand, high *CD44v2-10, v3-10, and v8-10* were characteristic of the basal-like group, whereas *CD44*v10 and *CD44*s were characteristic of the EMT phenotype, reinforcing the close link between shorter isoforms resulting from mRNA processing and invasive traits. Notably, in this cohort, the three subgroups presented similar survivals explained by the aggressive nature of the tumors. Nevertheless, we showed the existence of different molecular subtypes for tumors of apparently similar histology and outcome, including CD44 signatures that should be confirmed foreseeing precise cancer targeting.

Collectively, transcriptomics strongly supports the co-existence of multiple CD44 proteoforms in the healthy urothelium and cancer. It also highlights the close link between the presence of shorter isoforms resulting from alternative splicing and muscle-invasive disease, characterized by unfavorable outcomes. Nevertheless, confirmation by high-throughput proteomics is required for definitive confirmation.

### CD44s variant is expressed in healthy human tissues

Precise targeting requires significant cancer specificity. Therefore, we also evaluated a wide number of healthy tissues that exhibited variable but always detectable *CD44* mRNA expression (**Fig. 1E**). This highlighted the complex mosaicism presented by CD44 in human organs, later confirmed by consultation of the Genotype-Tissue Expression (GTEx) database (**SI Appendix, Fig. S3**). In addition, RT-PCR (**Fig. 1E**) and whole transcriptome analysis (**SI Appendix, Fig. S3**) show that, despite overexpressed in bladder cancer, CD44s may also be strongly expressed by most healthy cells, which significantly challenges its cancer specificity. Therefore, addressing CD44s post-translational modifications, namely glycosylation, poses as the next logical step towards this objective.

### BC expresses CD44 glycoproteoforms not observed in relevant healthy cells/organs

In the past, we and other groups have demonstrated how glycosylation may increase the cancer specificity of proteins, which could be explored for precise cancer-targeting (25, 37-41). CD44 presents multiple potential *O*-glycosylation sites in its extracellular domain, mostly in the variable region, which could be explored towards this objective (2). We hypothesized that CD44 may reflect the profound alterations occurring in glycosylation pathways in BC, which have been directly linked to aggressive traits. As such, we have screened BC sections of different histological natures by immunohistochemistry for signs of co-localization between CD44 and the Sialyl-Tn (STn) antigen, an immature *O*-glycan promoter of invasion (42, 43), immune escape (44), and an independent predictor of poor prognosis (45). We have also studied the Tn antigen, the shortest *O*-glycan, whose expression in BC has been linked to cancer aggressiveness in explorative studies. To avoid any possible cross-reactivity between the VVA lectin used to detect the Tn antigen and blood group A antigens, tissue sections positive for the A antigen were excluded from the analysis. According to **Figs. 2A-B**, neither the Tn nor the STn could be observed in healthy urothelium, except for some Tn positivity in the cytoplasm of upper stratum umbrella cells. STn antigen levels were significantly elevated in tumors in comparison to the healthy urothelium, being statistically significant for invasive lesions (**Fig. 2A**), as previously reported by us (36, 45-47). It was mostly detected at the surface of cancer cells in both superficial and invasive tumor layers, being more intense in invasive fronts in accordance with its functional role in disease progression (36, 39). The less studied Tn antigen was not significantly overexpressed in bladder tumors, except for a subgroup of patients at more advanced stages. In BC, the Tn antigen was predominantly cytoplasmic, most likely due to the detection of immature glycans in protein secretory pathways. For 47% of the cases (data not shown), membrane expression was also evident, but no pattern could be established regarding location, proximity to vessels or any other histopathological characteristics. Nevertheless, all tumors exhibiting abnormal glycosylation, also presented areas of CD44-Tn and/or STn co-expression (**Fig. 2B**), irrespectively of their histological nature. Also, *in situ* proximity ligation assays (PLA; **Fig. 2C**) and double staining immunofluorescence (**Fig. 2D**) for CD44 and STn and Tn antigens, respectively, revealed close spatial proximity between CD44 and the glycans in tumors but not in the healthy urothelium. Notably, positive PLA for CD44-STn was mainly observed in invasive fronts (**Fig. 2C**) consistent with the role played individually by CD44 and STn in cancer invasion (11, 48).

**Fig. 2.**
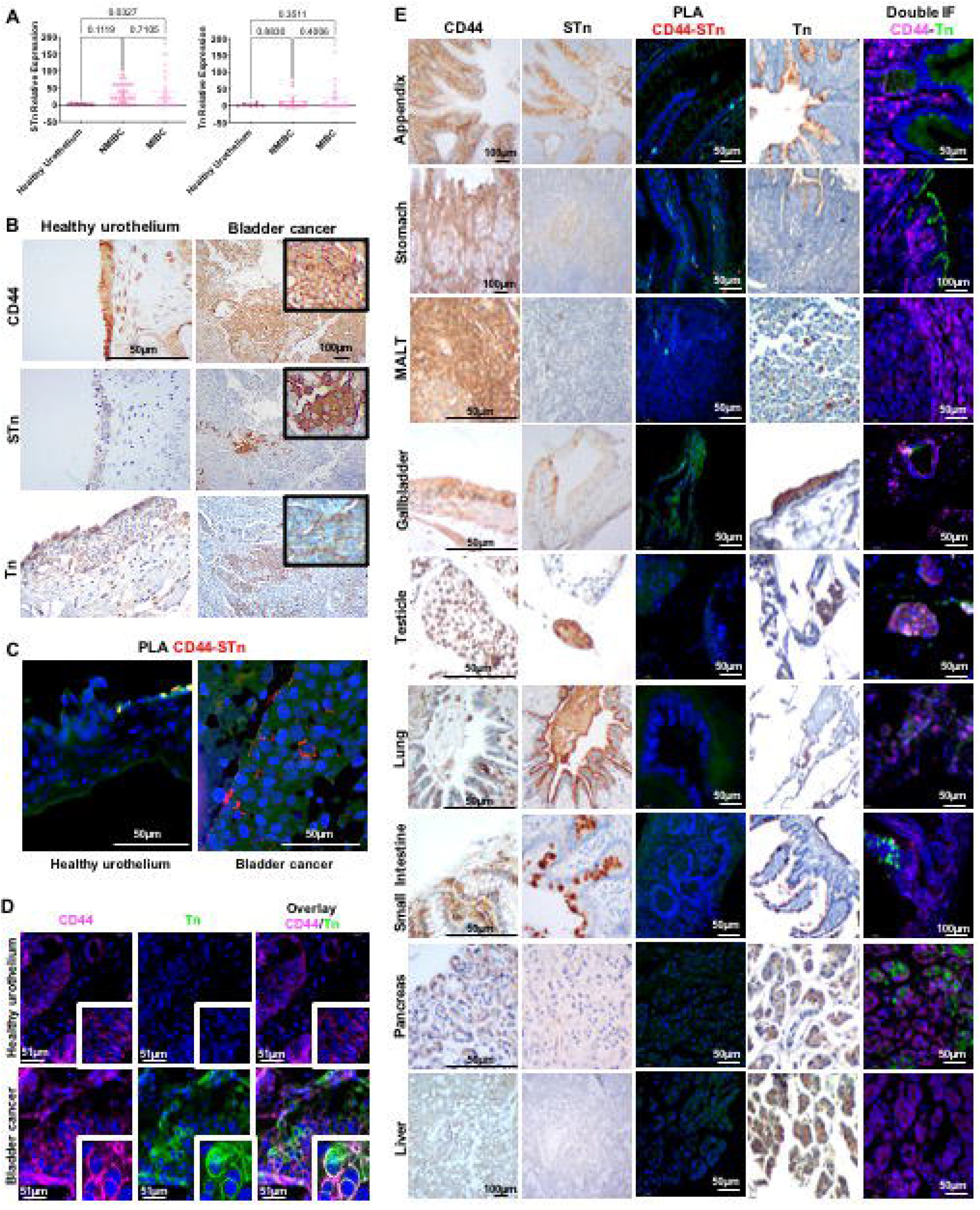
CD44-Tn and STn glycoproteoforms present high cancer specificity. **A) Tn and STn antigens are not expressed in the healthy urothelium, are elevated in cancer, and increased in invasive tumors**. The STn antigen, defined by affinity for the anti-tag-72 antibody [B72.3+CC49], is not expressed in the healthy urothelium and is elevated in cancer, being significantly overexpressed in muscle invasive bladder cancer (MIBC). The Tn antigen, based on VVA lectin immunoaffinity, is also not expressed in the healthy urothelium and is elevated in a subset of patients, specially at more advanced stages. **B) CD44s colocalizes with Tn and STn antigens in MIBC**. CD44 was diffusively expressed across the tumor section without a defined pattern. The STn antigen was observed both in superficial and invasive layers and the Tn antigen was found in scattered niches without a defined expression pattern. Immunohistochemistry also showed the co-localization of CD44 with Tn and STn positive areas in CD44s^high^ tumors, exhibiting low amounts of other isoforms (according to RT-PCR, data not shown). The healthy urothelium expressed high amounts of CD44 and the Tn antigen is present in the cytoplasm of upper stratum umbrella cells while the STn antigen was not detected. **C) *In situ* proximity ligation assays (PLA) supports CD44s-STn glycoproteoforms in MIBC and its presence within tumor invasive fronts**. *In situ* proximity ligation assay showed close spatial proximity between CD44 and STn in the same cells, strongly supporting CD44-STn glycoproteoforms in tumors. This phenotype was mostly observed in invasive fronts of CD44s^high^ tumors and was not detected in the healthy urothelium, suggesting cancer-specificity. **D) Double staining immunofluorescence supports CD44s-Tn glycoproteoforms in bladder cancer**. CD44s^high^ tumors presented niches of cells co-expressing CD44 and Tn antigen, strongly suggesting CD44-Tn glycoproteoforms. This was not observed in the healthy urothelium. **E) Glycosylation with Tn and STn antigens provides cancer specificity to CD44**. CD44 was abundantly expressed in all studied healthy tissues. STn expression was either absent or low in secretions and cells facing the lumen of the respiratory, gastrointestinal, and colorectal tracts. Immunohistochemistry suggested STn and CD44 co-localization in the stomach, appendix, small intestine, colon, gallbladder, and white blood cells in mucosa-associated lymphoid tissue (MALT). PLA did not confirm these hypotheses, suggesting cancer specificity of CD44-STn glycoproteoforms. In healthy tissues, the Tn was restricted to the cytoplasm of goblet cells in the intestinal tract, Leydig cells in testicular tissue, pancreatic acini, hepatocytes, mucinous cells of the gastric epithelium, alveolar macrophages, and gallbladder epithelium. CD44 and Tn antigen co-expression was suggested in pancreatic tissue, testicle, and gallbladder, which was not confirmed by double staining immunofluorescence.

We have further investigated the cancer specificity of the CD44-Tn/STn proteoforms in healthy tissues. STn expression was either absent or low in secretions and cells facing the lumen of the respiratory, gastrointestinal, and colorectal tracts (**Fig. 2E**), in accordance with our previous reports (39, 41). Amongst STn-positive tissues, we found possible co-localization between the glycan and CD44 in the stomach, appendix, small intestine, colon, the gallbladder, and white blood cells in mucosa-associated lymphoid tissue (MALT; **Fig. 2E**) and in peripheral blood mononuclear cells from healthy donors (data not shown). Interestingly, these organs also showed higher levels of cancer associated CD44s. However, orthogonal validation by double staining immunofluorescence and PLA did not confirm possible CD44-STn glycoproteoforms in healthy tissues, suggesting cancer specificity (**Fig. 2E)**. On the other hand, the Tn antigen was circumscribed to the cytoplasm of goblet cells at the intestinal tract, Leydig cells at testicle, pancreatic acini, hepatocytes, mucinous cells at gastric epithelium, alveolar macrophages at lung and gallbladder epithelium. Immunohistochemistry suggested some degree of overlap between Tn and CD44 in the pancreas, testicle, and gallbladder, which showed low CD44s expression (**Fig. 2E**). However, the presence of CD44-Tn glycoproteoforms was not supported by immunofluorescence (**Fig. 2E**). Collectively, according to our observations, short-chain *O*-glycans provide cancer-specificity to CD44 and the means for precise cancer targeting. The high abundance of CD44s compared to other isoforms in tumor tissues strongly supports this isoform as a major carrier of altered glycosylation.

### Glycoproteogenomics identifies multiple glycoproteoforms in BC cells

Transcriptomics and different immunoassays strongly supported the existence of BC specific CD44 glycoproteoforms linked to different stages of the disease, which require precise identification by proteomics. Therefore, we started by choosing cell models to support the implementation of a mass-spectrometry based roadmap to address this challenge. Three well established BC cell lines reflecting distinct stages of the disease (derived from a grade I (RT4), a grade II (5637) and grade III (T24) carcinomas) were evaluated for the capability to invade Matrigel *in vitro* and screened for CD44 by Real Time-Polymerase Chain Reaction (RT-PCR), flow cytometry, and western blotting. CD44 expression increased alongside with cell lines invasive capacity (T24>5637>RT4; **Figs. 3A-B)**, in accordance with the histopathological nature of the carcinomas from which cell lines derived. We also found that less invasive RT4 and 5637 cells expressed lower levels of CD44 in comparison to T24 cells (**Figs. 3B-C**), being also enriched for longer CD44 variants (**Figs. 3D-E**). On the other hand, T24 cells expressed high levels of CD44s and residual amounts of other variants (**Figs. 3B-E**), in agreement with patient samples observations linking this variant to invasive traits. Therefore, we considered T24 cell line as a suitable model to study the influence of CD44s in BC, since the other variants are practically vestigial. Moreover, based on the close similarity between RT4 and 5637, we selected 5637 and T24 cells for downstream proteomics-based studies.

**Fig. 3.**
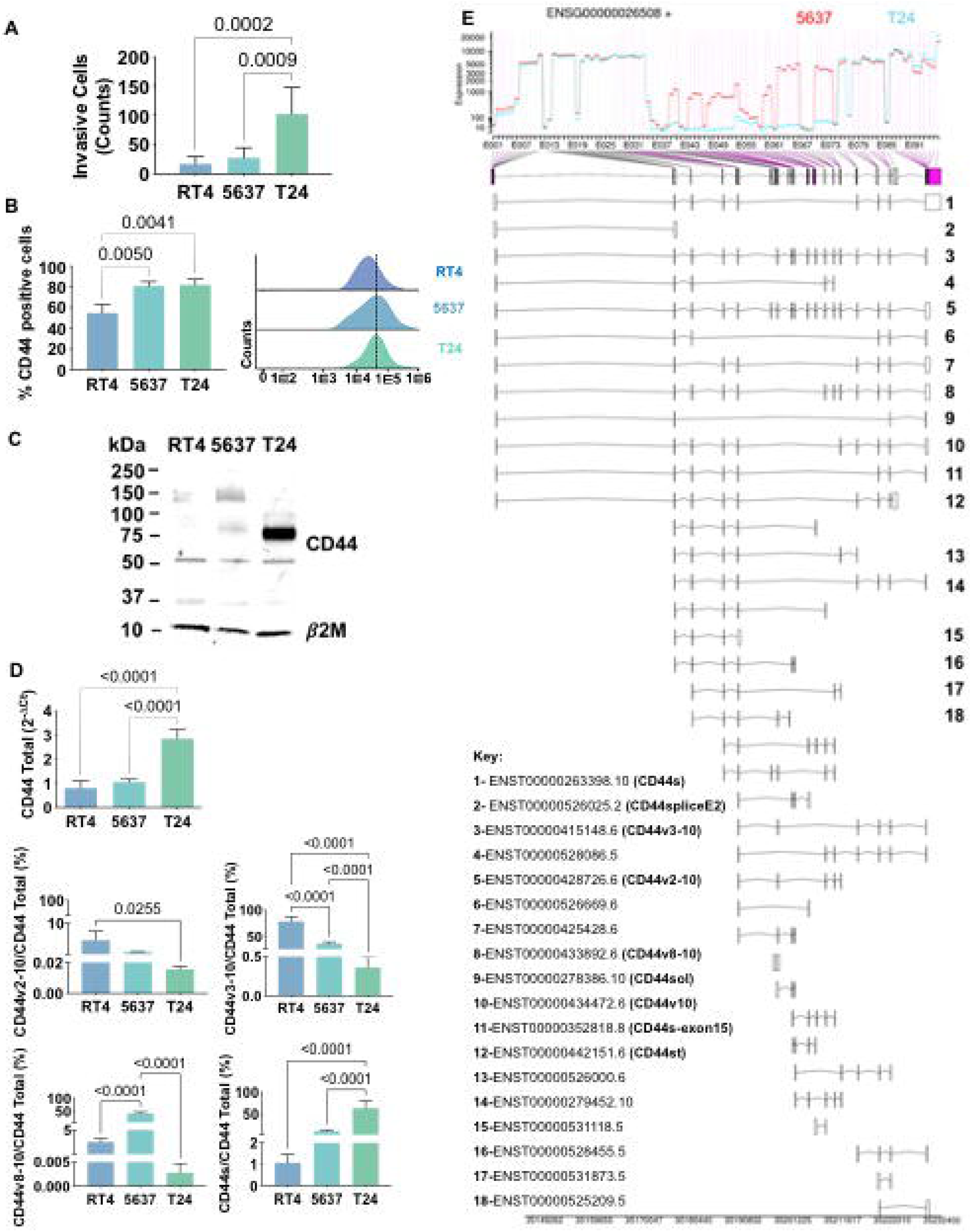
Highly invasive T24 bladder cancer cells express high levels of CD44s. **A) Capacity to invade Matrigel *in vitro* for RT4, 5637 and T24 cells**. Grades I/II cell lines (RT4 and 5637) are significantly less capable of invading Matrigel *in vitro*. **B) CD44 expression is higher for grade II (5637) and III (T24) cells**. Flow cytometry analysis showed significantly higher CD44 levels in 5637 and T24 compared to RT4 cells. **C) Western blots show different CD44 expression patterns according to cell grade, with T24 cells showing mostly shorter proteoforms**. Western blotting confirmed the overexpression of shorter CD44 proteoforms (at approximately 75 and 50 kDa) in T24 cells, and the presence of heavier proteoforms in the other cell lines (above 150 kDa). **D) Characterization of CD44 isoforms by RT-PCR showing isoforms shortening with cell lines aggressiveness and the marked CD44s**^**high**^ **phenotype of T24 cells**. *CD44* gene expression was significantly higher in T24 in comparison to RT4 and 5637 cells, in agreement with protein analysis. RT-PCR also revealed increasing mRNA shortening with cells aggressiveness. Accordingly, RT4 showed higher *CD44v2-10* and *CD44v3-10* expression in comparison to the other cell lines, 5637 predominantly presented CD44v8-10, whereas T24 cells mainly expressed CD44s. **E) RNAseq confirmed the marked difference between T24 and 5637 cells, the first expressing shorter CD44 mRNAs in opposition to lengthier mRNAs in 5637 cells**. This result clearly highlights the significant splicing of *CD44* mRNA in T24 cells, originating isoforms missing the extracellular domain encoded by exons 6-14. The results correspond to the mean and standard deviation for three independent experiments. Triplicate measurements were conducted for each experiment. P values are presented for one-way ANOVA tests.

In explorative studies using conventional bottom-up proteomics approaches, we concluded that the precise characterization of CD44 glycoproteoforms could only be accomplished by the customization of the databases used for protein annotation and by upfront knowledge of the cellular glycome. Through transcriptome analysis of 5637 and T24 cell lines, we discovered 38 different RNA transcripts of CD44, 18 of which have been previously described (**Figs. 3D-E; SI Appendix, Table S2**). Amongst these, 8 transcripts had already been experimentally validated (including CD44v2-10, CD44v3-10, CD44v8-10 and CD44s herein addressed by RT-PCR; **Fig. 3D**) and 3 were related to non-coding RNA (**Fig. 3E; SI Appendix, Table S2**). The remaining 7 transcripts correspond to computationally inferred short transcripts (**Fig. 3E; SI Appendix, Table S2**). However, half lacked open-reading frames (ENST00000526000.6 [Uniprot:H0YDW7]; ENST00000279452.10 [Uniprot:H0Y2P0]; ENST00000528455.5 [Uniprot:H0YD17]; ENST00000531873.5 [Uniprot:H0YD90]) are unlikely to be translated into proteins, while another contained a premature translation-termination codon (ENST00000425428.6 [Uniprot:Q86UZ1]). Transcript ENST00000526669.6 (Uniprot:H0YD13) encoded an incomplete extracellular domain, lacked membrane anchoring and/or intracellular region, and presented low transcript levels, suggesting reduced probability of occurrence. Finally, transcript ENST00000442151.6 (Uniprot:H0Y5E4) presents almost 100% homology with CD44st with the exception of a minor 5’truncation corresponding to the first amino acid in the protein sequence. In summary, only the computationally determined variant ENST00000442151.6 was considered for downstream validation by mass spectrometry. The remaining 20 transcripts corresponded to previously undescribed RNA sequences, probability corresponding to short non-coding transcripts, whose biological role should be investigated in the future. Collectively, transcriptomics confirmed the intense alternative splicing of CD44 in T24 cells, leading to several short transcripts encoding proteins lacking a variable extracellular region. After excluding ambiguous transcripts, a total of 9 sequences were considered plausible to generate proteins and were selected to construct the CD44 protein database (**SI Appendix, Table S2**).

In addition, we have characterized the 5637 and T24 cells *O*-glycome by MALDI-MS. Adding to distinct clinicopathological features, these cells exhibited slightly different *O*-glycomes in terms of glycan abundance (**Fig. 4A**). Both cells predominantly expressed core 1 derived structures, namely fucosyl-T (*m/z* 768.38; [M+Na]^+^) and mono- (*m/z* 955.46) and di-sialylated T antigens (*m/z* 1316.64) (**Fig. 4A**). However, in 5637, fucosylated and mono-sialylated T antigens were more abundant, whereas in T24 it was disialyl-T. The presence of several extended core 2 glycans (*m/z* 1217.60; m/z 1391.69; 1404.69; 1578.78; 1765.86) were also observed, being however more pronounced in 5637 cells. Low amounts of core 3 (*m/z* 635.32) and T (*m/z* 594.29) antigens were found in both cells as well as STn antigens (*m/z* 751.36) in T24. Overall, key information was generated to guide CD44 glycosites identification, contributing to increase protein coverage in glycoproteogenomics settings.

**Fig. 4.**
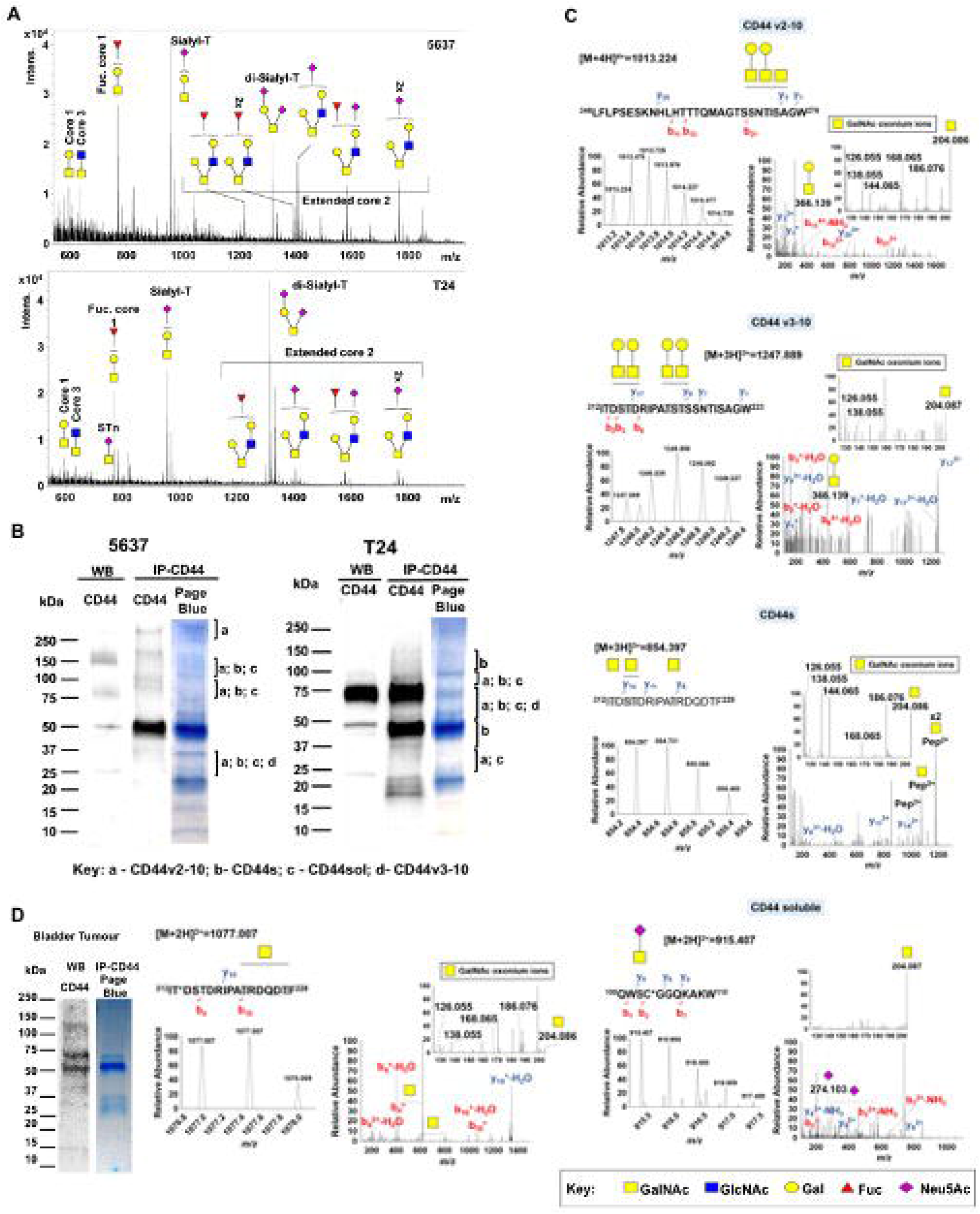
Glycoproteogenomics, building on RNAseq-customized databases and glycomics for protein annotation, enables CD44 proteoforms identification in bladder cancer cell lines and tumors. **A) Glycomics characterization of 5637 and T24 cells showed fucosylated and sialylated T antigens as main glycospecies**. MALDI-MS analysis of permethylated benzyl-GalNAc glycans revealed [M+Na]^+^ main ions for fucosylated and sialylated T antigens in both cell lines. However, 5637 predominantly expressed mono-sialylated T antigens whereas di-sialylated structures were more abundant in T24 cells. Moreover, 5637 cells presented higher abundance and diversity of glycoforms extended beyond core 2 with different degrees of sialylation and fucosylation. **B) SDS-PAGE gels and western blots for CD44 IPs highlighting the nature of the isoforms identified by nano-LC-M/MS in glycoproteogenomics settings**. Briefly, CD44 immunoprecipitated from membrane extracts was separated by gradient SDS-PAGE and bands were excised and analyzed by nanoLC-HCD/CID-MS/MS using RNAseq-customized databases and glycomics data for isoforms annotation. The nature of the isoforms identified in each band and cell line was highlighted. CD44 heterogeneity was evident for the two cell lines. CD44s was found to be the main isoform in T24 cell line. **C) nano-LC-MS and HCD-MS/MS spectra for reporter ions of CD44v2-10, CD44v3-10, CD44s, and CD44sol in T24 cells**. For the reporter ions we show the MS isotopic envelope, the corresponding HCD product ion spectra highlighting GalNAc oxonium ions, GalNAc cross-ring fragments, other glycan fragments, and y- and b-series peptide backbone ions that support protein annotation. The predicted glycopeptide sequence, including the nature of the glycans and glycosites annotation (whenever possible) was also presented. **D) nano-LC-MS and HCD-MS/MS spectra for CD44 immunoprecipitated from CD44s**^**high**^ **tumors and areas of CD44-STn co-expression**. SDS-PAGE and western blots showed a pattern similar to T24 cells, characterized by bands bellow 75 kDa. An HCD product ion spectrum for an CD44s-Tn specific glycopeptide sequence is presented. Identified and possible glycosites are identified in grey. The symbol * corresponds to amino acid modifications: C – carbamidomethyl and T – 2-amino-3-ketobutyric acid.

For glycoproteoforms identification (**SI Appendix, Fig. S4)** we first immunoprecipitated CD44 from plasma membrane-enriched protein extracts isolated by differential ultracentrifugation. To ensure broad CD44 representation, we adopted an antibody targeting the cytoplasmatic tail, which is conserved amongst the different known splice variants with the exception of CD44st. This avoided glycosylated regions of the protein, which may experience significant structural variations due to the dynamic nature of this post-translational modification, thus affecting antibody recognition. We first separated different proteoforms by gradient SDS-PAGE supported by western blot, excised the bands from gels (**Fig. 4B**) and applied a proteomics workflow contemplating digestion with chymotrypsin and protein identification by nanoLC-HCD-MS/MS. This approach was complemented by CID-MS/MS triggered by the presence of the oxonium ion HexNAc (*m/z* 204.087) in the HCD-MS/MS. To render the glycosylation more homogeneous and facilitate downstream assignments by MS/MS, the glycoproteins were desialylated prior to proteolytic digestion. The western blots in **Fig. 4B** started by highlighting the diversity of proteoforms across a wide range of molecular weights in BC cell lines **(Figs. 4C-D)**, as suggested by transcripts analysis. Western blots also reflected the abundance of CD44 in the cell lines, previously suggested by RT-PCR and later confirmed by flow cytometry. CD44 molecules bigger than 150 kDa predominated in 5637 cells, whereas T24 cells presented a major band at 75 kDa. Interestingly, both cell lines presented several well defined proteoforms below 37 kDa.

In explorative settings, we observed that unambiguous isoforms characterization was not possible without considering *O*-glycosylation, even when using transcriptome-customized databases. An integrative glycoproteogenomics approach was then applied, which enabled the identification of 4 isoforms (CD44v2-10, CD44v3-10, CD44s, CD44sol) with high degree of confidence, based on specific diagnostic glycopeptides (**Figs. 4B-C**; **SI Appendix, Tables S3 and S4; SI Appendix, Fig. S5**). Using HCD fragmentation, we could identify several product ions consistent with the presence of glycosylated moieties, together with some *y-* and *b*-type peptide fragments supporting these reporter glycopeptides, even though with some ambiguity in terms of glycosites assignment (**Fig. 4C**). The presence of GalNAc, the first sugar residue in *O*-glycans, could be further confirmed by several HexNAc cross ring fragments (*m/z* 126.055,168.066,186.076, and 204.087; **Figs. 4C-D** and **SI Appendix, Figs. S5-6 and 10**) and, in some cases, high 144.066/138.055 oxonium ions ratios (49, 50). We also explored HexNAc-triggered CID fragmentations for further glycopeptide validation, including the characterization of extended glycan chains (**SI Appendix, Fig. S5)**. For certain species, CID-based fragmentation retrieved a significant number of peptide backbone fragmentations, enabling unequivocal glycosites characterization, thus in accordance with our previous reports using a similar approach (51). Notably, we found CD44 glycoproteoforms carrying both short- and elongated glycans, as well as the co-existence of different glycoforms in the same peptide sequence, even though in distinct glycosites (**Fig. 4C**; **SI Appendix, Tables S3 and S4; SI Appendix, Fig. S5)**. Also, despite subjected to desialylation, some glycopeptides carrying sialic acids were still observed due to incomplete enzymatic digestion (**Fig. 4C; SI Appendix, Fig. S5**). Nevertheless, such observations, reinforce the tremendous molecular micro-, macro-, and meta-heterogeneity presented by CD44, previously reported for other human glycoproteins (24), portraying the relevance of glycosylation towards unequivocal molecular characterization.

Overall, for 5637 cells, we identified high molecular weight CD44v2-10 at 250 kDa, as well as CD44v2-10, CD44s and CD44sol in several bands above 75kDa, most likely resulting from heterogeneous glycosylation and multiple glycosites occupation (**Fig. 4B; SI Appendix, Table S3**). An intense band at 50 kDa was also present corresponding to the anti-CD44 antibody used for IP, since no CD44 isoforms were identified at this molecular weight. In T24 cells, CD44s was identified in bands spanning from 150 to 50 kDa, including major bands at 75 and 50 kDa, in agreement with its higher abundance in these cells (**Fig. 4B, SI Appendix, Table S4**). In addition, we could also identify the other three isoforms above 75 kDa, again reinforcing CD44 heterogeneity. Interestingly, both cell lines presented low molecular weight proteoforms below 37 kDa corresponding to CD44v2-10 and CD44v3-10, whose full-length proteins are generally present above 80 kDa (without considering glycosylation). We believe these may be products of proteolysis occurring in extracellular domains that remained anchored to the cell membrane, which warrants confirmation. On the other hand, the CD44v8-10 variant observed by RT-PCR and transcriptomics (**Fig. 3D**) could not be confirmed by mass spectrometry. Interestingly, our glycoproteogenomics approach identified CD44v2-10, v3-10 and CD44sol, which showed far lower transcripts levels in comparison to CD44v10 (**Fig. 3D**). Even though missing definitive confirmation, these observations reinforce the gaps between the transcriptome and the proteome and the importance of exploring MS-based methodologies for precise proteoforms characterization. In summary, we have shown that CD44 characterization cannot be achieved by a single omics, requiring glycoproteogenomics settings. Moreover, addressing glycosylated domains is key for precise CD44 characterization.

### Glycoproteogenomics identified cancer specific CD44s glycoproteoforms in bladder tumors

Transcriptomics and different immunoassays strongly suggested but cannot present unequivocal demonstrations regarding the existence of CD44s-STn/Tn glycoproteoforms in BC. To address this limitation, our glycoproteogenomics approach was further employed to interrogate tumors for short glycoproteoforms specifically linked to BC aggressiveness. A protein pool from invasive tumors of five different patients showing high CD44s transcripts, CD44 western blot patterns resembling T24 cells (**Fig. 4D**), and high STn and Tn antigens expressions was elected for this study. Immunofluorescence assays suggesting the presence of CD44-STn and Tn glycoproteoforms in the same tumor area led to consider these glycans during protein annotation. CD44-STn/Tn co-expressing areas were then excised from paraffin-embedded tumor sections after deparaffination, antigen retrieval, proteins were then extracted, and CD44 was isolated by immunoprecipitation and characterized as demonstrated for BC cell lines. Despite the significant degree of protein alterations induced by the conservation of the tumors during histological procedure, namely through multiple oxidations, we successfully identified CD44s in tumors (**Fig. 4D**). Furthermore, we could identify the presence of short-chain cancer-associated *O*-glycans in both the CD44 constant region (**SI Appendix, Fig. S5**) as well as in CD44s reporter peptide sequences (**Fig. 4D)**. This constitutes unequivocal proof of aberrant CD44s glycosylation in BC, confirming immunoassays.

### CD44s is a driver of invasion

We then investigated the functional role played by CD44 in proliferation and invasion by silencing its expression in 5637 and T24 cells using siRNA (80-90% CD44 decrease according to RT-PCR and western blot; **Figs. 5A-B**). This significantly reduced cell proliferation in both cell lines (**Fig. 5C**) but impacted differently in terms of invasion, potentiating 5637 and decreasing the number of T24 invasive cells (**Fig. 5D**). These are novel findings linking CD44 to cell proliferation in BC, supporting previous observations for breast (52) and colorectal (53) cancers, and suggesting that intense splicing of extracellular domains may play a key role in driving invasion, in agreement with observations from patient samples.

**Fig. 5.**
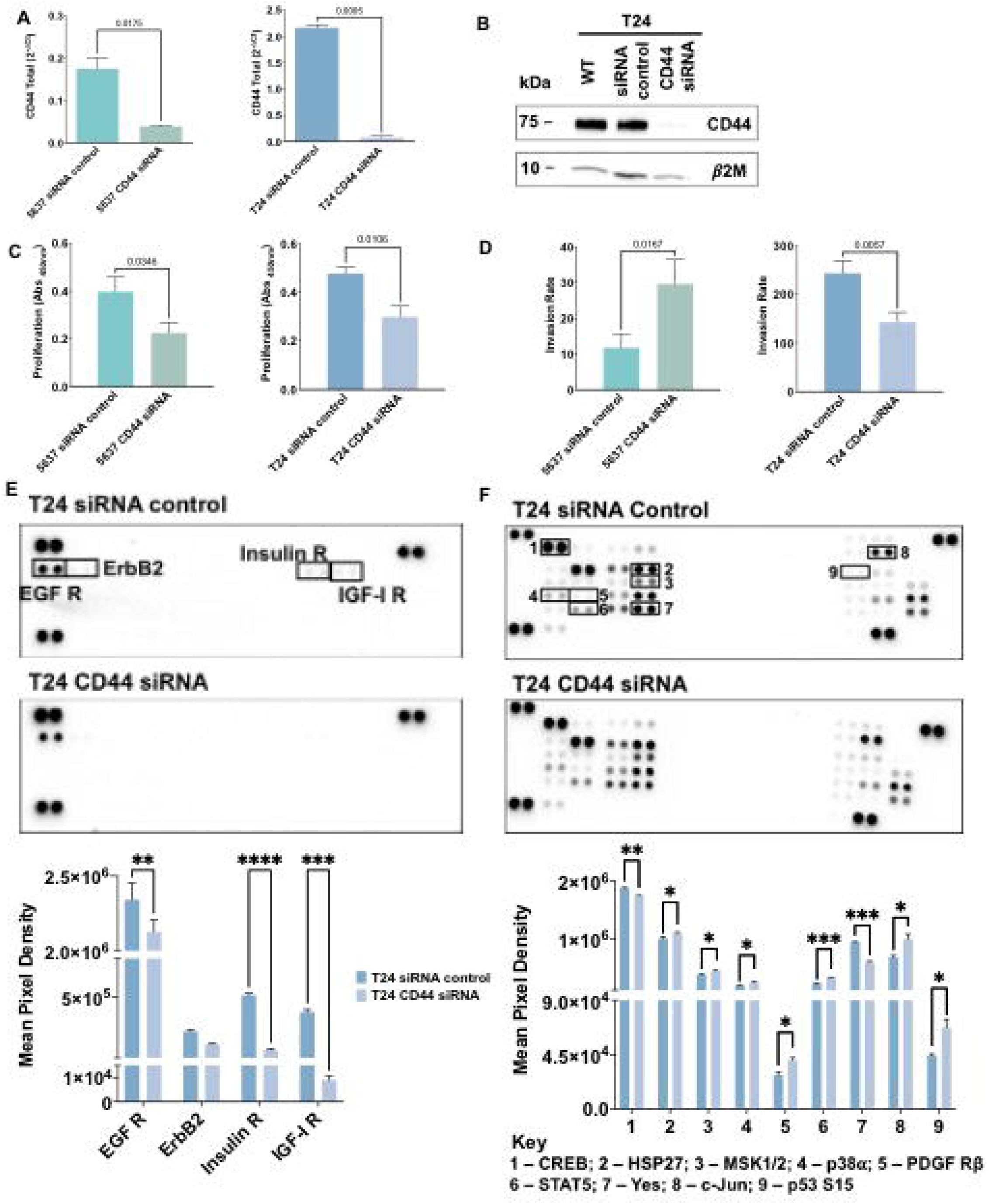
CD44 silencing inhibits proliferation, invasion, and activation of relevant proteins in MAPK/ERK, and PI3K/AKT pathways in T24 CD44s^high^ cells. **A and B) CD44 siRNA promotes a massive decrease in *CD44* gene expression and complete abrogation of CD44 protein levels in 5637 and T24 cell lines**. According to Figure A, *CD44* gene expression after siRNA is decreased by approximately 80% and 96% in 5637 and T24 cells, respectively, in comparison to controls, which translated into significantly decreased protein expression (exemplified in Figure B for T24 cells). **C) CD44 is a promoter of proliferation in 5637 and T24 cells**. Proliferation was reduced by more than 50% after CD44 siRNA silencing in both cell lines compared to controls. **D) CD44 silencing decreases the invasive capacity of CD44s**^**high**^ **T24 cells and promoted invasion in 5637 cells**. The number of T24 invading cells decreased approximately 1.7-fold after CD44 knock-down comparing to controls. On the other hand, the number of cells invading matrigel *in vitro* was 1.7-fold higher for 5637 cells under the same stimuli. **E) Analysis of the phosphorylation status of 49 receptors tyrosine kinases (RTK) and F) Effect of CD44 silencing in phosphorylation of 37 proteins activated by different kinases**. CD44s seems to be a modulator and a co-receptor of different RTKs, including EGFR, Insulin, and IGF-1 receptors, as well as transcription factors such CREB, involved in tumor proliferation and invasion. The results correspond to mean and standard deviation for three independent experiments. Triplicate measurements were conducted for each experiment. P values are presented for Unpaired T tests.

To disclose how CD44 could mediate proliferation and invasion, we used a human Phospho-Receptor Tyrosine Kinase (Phospho-RTK) array to detect possible alterations in the phosphorylation levels of 49 different proteins **(SI Appendix, Fig. S7)** from T24 cells after CD44 siRNA-mediated knockdown. EGFR, a cancer-associated member of the ErbB family, was significantly less activated in CD44 silenced T24 cells when compared to controls, which is consistent with CD44/EGFR interaction and synergic oncogenic signaling towards proliferation and invasion (54, 55). Similar behavior was observed for the highly homologous insulin (IR) and insulin-like growth factor 1 (IGF-1R) receptors (56) after CD44 silencing **(Fig. 5E)**. Interestingly, increased IGF-1R phosphorylation in CD44 expressing cells has been linked to concomitant PI3K/AKT/mTOR pathway activation, EMT phenotypes and stem cells maintenance in solid tumors (57). These observations support that CD44-mediated IR and IGF-1R activation, including downstream AKT and MAPK signaling transduction governing proliferation and invasion (58, 59) may also occur in BC. Altogether, CD44s, the most prevalent variant in T24 cells, seems to promote the activity of relevant RTK, culminating in biological cues that drive cancer cell proliferation and invasion. However, the exact mechanism by which CD44 triggers RTK activation remains poorly understood and should be addressed in future studies.

We further investigated how CD44 silencing would influence the activation by phosphorylation of several phospho-kinases and related downstream targets, using a human Phospho-Kinase array **(SI Appendix, Fig. S8)**. As shown in **Fig. 5F**, the activating phosphorylation of the transcription factor CREB as well as Src family non-receptor tyrosine kinase Yes was significantly decreased in T24 cells upon CD44 knock-down. CREB has been previously reported as a downstream target of CD44, mediating tumor proliferation, progression, and invasion in several tumor models (60, 61). CD44 knock-down also resulted in HSP27, MSK1/2, p38α, PDGFRβ, c-Jun, STAT5a/b, Yes and p53 (S15) phospho-kinase activation. In agreement with these observations, several studies have described CD44 as a negative regulator of PDGFRβ signaling by mediating the recruitment of a tyrosine phosphatase to PDGFR and by destabilizing PDGFR heteroreceptor complexes, ultimately modulating growth factor signaling and proliferation (62, 63). Moreover, CD44 has been previously described as a negative regulator of p53 mediated responses, allowing cells to resist programmed death and senescence as well as respond to proliferative signals (64, 65). The regulatory effect of CD44 on HSP27, MSK1/2, STAT5a/b and Yes are being described for the first time and warrant deeper investigation. Notably, Yes is the most widely expressed member of the Src kinase family and has a known regulatory role in growth factor signaling, cell proliferation, and invasion in different cancer models (66, 67). Our observations support that CD44 may act as a Yes regulator in BC, which also merits future validation. On the other hand, CD44 has been described as a positive regulator of p38α (68) and c-Jun (69) in other tumor models, which potentially conflicts with our current observations and suggests cell-dependent responses that may be governed by the nature of CD44 isoforms. Collectively, our observations emphasize that we are still far from fully understanding CD44 regulation of oncogenic pathways, which may now be comprehensively addressed in the context of its isoforms. Nevertheless, we provide evidence that CD44s acts as an oncogenic modulator of phospho-kinases involved in several signaling pathways that support proliferation and invasion, including ERK/MAPK, PI3K/AKT signaling pathways.

### CD44s glycosylation modulates proliferation and invasion

To assess the functional impact of altered CD44 glycosylation, we explored glycoengineered T24 cells overexpressing Tn (T24 *C1GALT1* knock-out (KO)) and STn antigens (T24 *C1GALT1*KO/*ST6GALNAC1* knock-in (KI)) in mimicry of aggressive bladder tumors. Tn antigen expression was accomplished by complete abrogation of *O*-glycans extension by *C1GALT1* KO, as translated by a complete loss of sialylated T antigens (**Fig. 6A**). Alongside the Tn antigen, these cells also expressed low amounts of STn, denoting low levels of sialylation in comparison to wild type cells. Then, *ST6GALNAC1* KI was introduced to promote Tn sialylation, generating a cell line expressing high STn levels and, to less extent, Tn antigens (**Fig. 6A**). According to immunofluorescence, all models co-expressed CD44 together with altered glycosylation at the cell surface (**Fig. 6A**). Furthermore, changes in glycosylation did not alter *CD44* gene expression or the nature of the variants, except for CD44s, which was found increased in cells overexpressing Tn but not for cells overexpressing STn (**Fig. 6B**). Western blots supported little changes in CD44 levels in glycoengineered models, as suggested by gene expression. However, the blots for glycoengineered cell models showed a slight decrease in the main molecular band at 75 kDa, most likely from changes in glycans complexity (**Fig. 6C**). This hypothesis was later confirmed by CD44 immunoprecipitation assays (**SI Appendix, Fig. S9**) and MS/MS analysis for T24 *C1GALT1KO* models (**Fig. 6D; SI Appendix, Fig. S10; SI Appendix, Table S6**). A less intense band above 37 kDa (close to CD44s estimated molecular weight) was also obvious in glycoengineered models (**Fig. 6C**), suggesting that *O*-glycans shortening may also impact on glycosites occupancy.

**Fig. 6.**
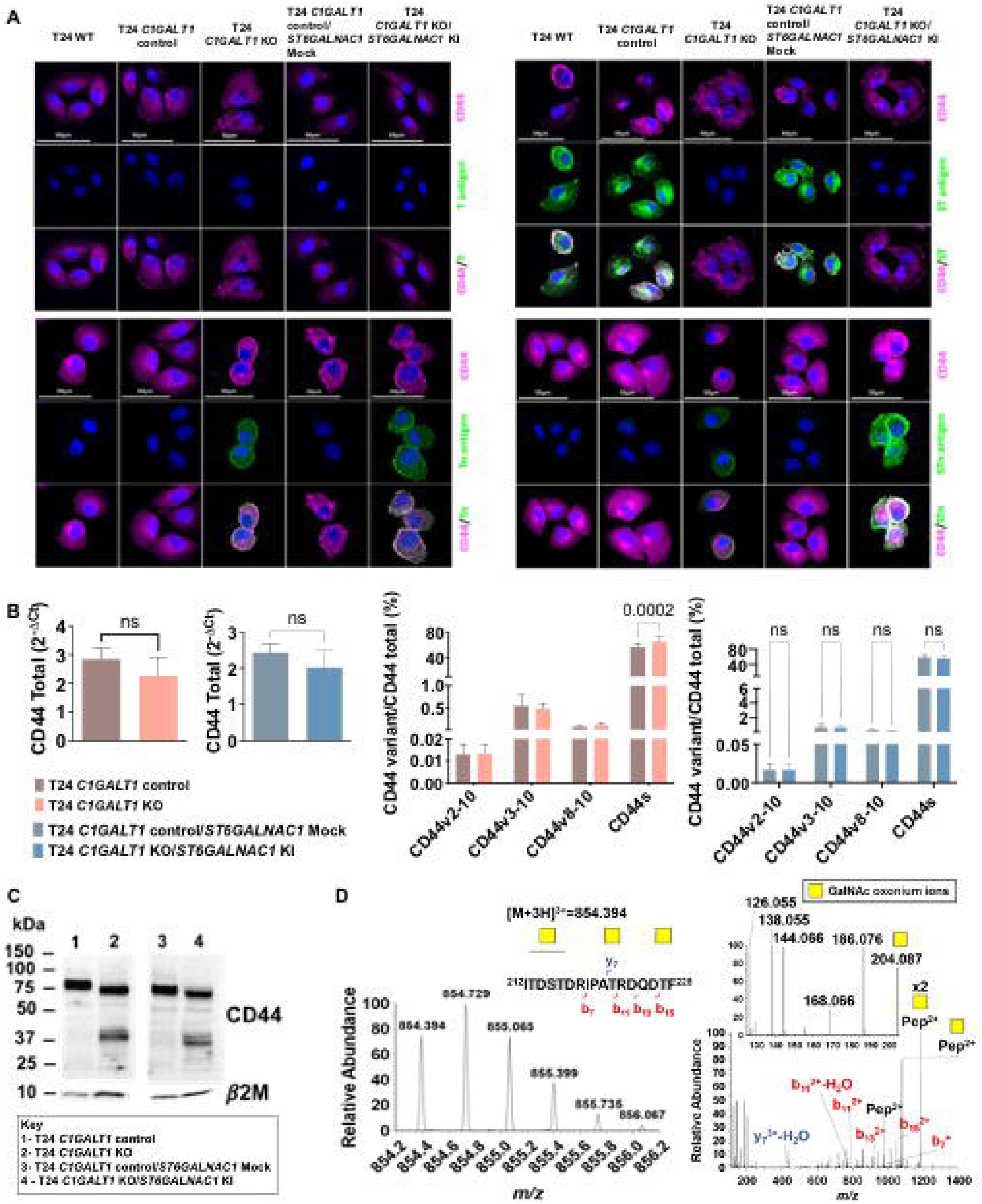
Glycoengineered T24 cells express CD44s-Tn/STn glycoproteoforms in mimicry of human tumors. **A) Immunofluorescence studies on glycoengineered T24 bladder cancer cells demonstrated the overexpression of immature short-chain *O*-glycans together with CD44 at the cell surface of the cells**. Immunoassays show high levels of sialylated T antigens and no Tn and STn antigens in wild type cells. T24 *C1GALT1*KOs lacked extended glycosylation translated by the presence of sialylated T antigens and presented high Tn levels. *ST6GALNA1*KI induced an overexpression of STn. These glycans were co-localized with CD44 at the cells surface. **B) *CD44* gene expression and the nature of the variants do not change with alterations in glycosylation induced by *C1GALT1*KO and *ST6GALNAC*1KI, except for *CD44s*, which increased in Tn-expressing cells. C) Western blots show a different CD44 expression pattern for T24 glycoengineered cells in comparison to controls**. The main molecular band of CD44 at 75kDa shows a slight decrease, possibly due to glycans complexity reduction. A new band at 37kDa emerged, suggesting that glycans truncation may impact on CD44 molecular weight. **D) MS and HCD-MS/MS for a CD44s specific glycopeptide from T24 *C1GALT1*KO cells evidencing short-chain *O*-glycosylation**. The product ion spectrum shows a typical GalNAc fragmentation pattern, characterized by the presence of an evident ion at *m/z* 144.066 and a high 144.066/138.055 oxonium ions ratio, as well as the presence of the others HexNAc oxonium ions (*m/z* 126.055,168.066, 186.076 and 204.087). The spectrum also shows major fragment glycopeptides with GalNAc losses. The results correspond to the mean and standard deviation for three independent experiments. Triplicate measurements were conducted for each experiment. P values are presented for two-way ANOVA and Unpaired T tests.

To disclose the influence of CD44 glycosylation, we induced a transient inhibition of the protein by siRNA in glycoengineered cell models and controls, which led to almost complete abrogation of *CD44* mRNA expression (**Fig. 7C**), as previously observed for wild type cells. First, we confirmed that proliferation was not affected by alterations in glycosylation induced by *C1GALT1*KO and *ST6GALNAC1*KI (**Fig. 7A**). Also, Tn overexpression did not change invasion, whereas STn-overexpression increased the number of T24 cells invading Matrigel (**Fig. 7B**). The link between STn and invasion is in close agreement with our previous report for glycoengineered BC cells (48), and other cancer models showing increased cell motility and acquisition of invasive traits resulting from the presence of this antigen (39, 42, 70). Nevertheless, subsequent studies showed that T24 *C1GALT1*KO and *C1GALT1*KO*/ST6GALNAC1*KI cells proliferation was not significantly affected by CD44s silencing (**Fig. 7D**). This contrasted with the anti-proliferative effect observed for T24 cells expressing more mature *O*-glycans under the same conditions (**Fig. 5C**). Such findings suggest that proliferation may be fine-tuned by the nature of CD44s *O*-glycosylation, being activated by *O*-glycans extension. On the other hand, invasion was decreased in STn-expressing cells after CD44 inhibition as previously observed for wild type cells (**Fig. 5D**). However, this effect was not observed for Tn-expressing cells, suggesting that sialylation may play a role in CD44-mediated invasion, which warrants future confirmation.

**Fig. 7.**
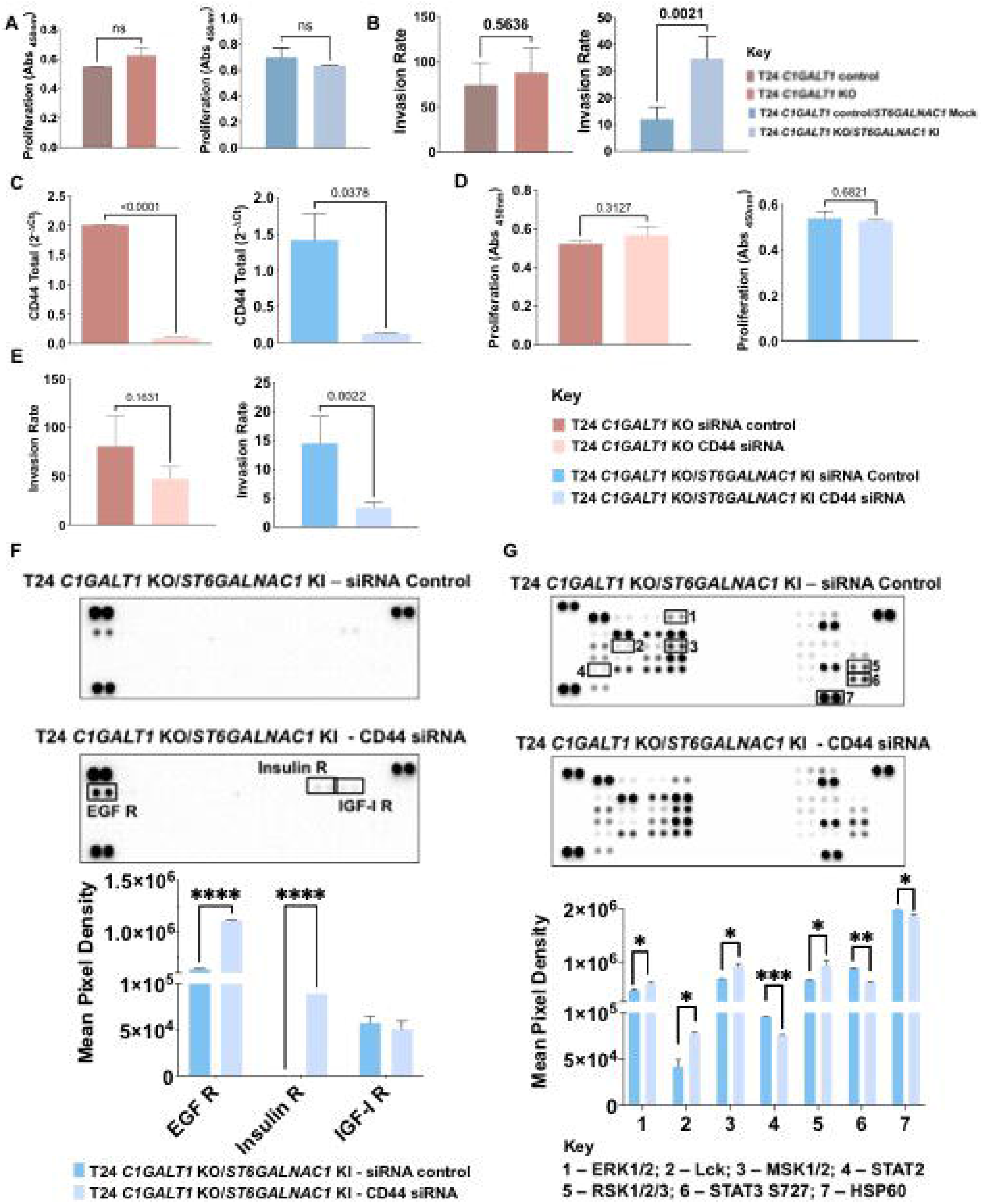
CD44s glycosylation shapes its functional contribution to proliferation and invasion. **A) *O*-glycans shortening by *C1GALT1*KO and *C1GALT1*KO/*ST6GALNAC*1KI does not change the proliferation of T24 cells**. T24 cells overexpressing Tn or STn antigens maintain proliferation rates compared to controls. **B) Increased STn antigen at the cell surface, driven by *C1GALT1*KO and *ST6GALNAC*1KI, enhances the invasive capacity of T24 cells**. T24 cells overexpressing the Tn antigen do not change their capacity to invade Matrigel in comparison to the control *in vitro*. On the other hand, STn overexpression leads to a significant increase in invasion. **C) CD44 silencing promotes a notable decrease in *CD44* gene expression in glycoengineered T24 cells expressing Tn and STn**. *CD44* mRNA expression after siRNA is significantly decreased by approximately 95% and 91% in *C1GALT1*KO and *C1GALT1*KO*/ST6GALNAC1*KI T24 cells, respectively, in comparison to controls. **D) CD44s-Tn and STn glycoproteoforms do not play a role in T24 cells proliferation**. After CD44s inhibition, *C1GALT1*KO and *C1GALT1*KO*/ST6GALNAC1*KI T24 cells maintained their proliferation in relation to controls, showing that CD44-Tn/STn glycoproteoforms do not influence cell proliferation. **E) CD44s-STn but not Tn glycoproteoforms drive invasion**. After CD44s silencing, *C1GALT1*KO T24 cells maintained their capacity to invade Matrigel *in vitro*, whereas *C1GALT1*KO*/ST6GALNAC1*KI cells showed decreased invasion. Similar effects were observed in wild-type cells but not in Tn-expressing cells, suggesting that sialylation may have a role in tumor invasion. **F) Influence of STn-overexpression on the response of tyrosine kinases receptors to CD44 silencing**. We show an activation of RTK proteins such EGFR and Insulin receptor in the absence of CD44 for *C1GALT1*KO*/ST6GALNAC1*KI cells, opposite to the effect observed in wild-type cells. **G) CD44s-STn glycoproteoforms seem to promote the activation of relevant transcription factors involved in cell proliferation and invasion**. In the absence of CD44, a significant downregulation of STAT3 S727 and STAT2 phosphorylation was observed in *C1GALT1*KO*/ST6GALNAC1*KI cells. Results correspond to the mean and standard deviation for three independent experiments. Triplicate measurements were conducted for each experiment. P values are presented for Unpaired T tests.

We have then attempted to understand the functional impact of STn-overexpression on the activation of different RTKs upon siRNA-mediated knockdown of CD44. As shown in **Fig. 7F**, CD44s silencing in STn positive cells induced a significant increase in activation of EGFR and Insulin receptors as well as of downstream proteins ERK 1/2, MSK 1/2 and RSK 1/2/3, all of which intermediates of the ERK/MAPK signaling pathway. Interestingly, wild type cells presented an opposite phenotype regarding activation of EGFR and insulin receptors under the same stimuli (**Fig. 5E**). These findings again portrait the key role played by the glycocode in the regulation of oncogenic pathways governed by CD44s.It further suggests that CD44s-STn driven invasion may be supported by other molecular mechanisms. Additionally, we showed increased phosphorylation of Lck, a member of Src kinase family, which is intimately involved in T-cell activation in the absence of CD44 (71), suggesting a broader role of CD44s in immune regulation that should be addressed in future studies. Furthermore, we observed a significant downregulation of STAT3, STAT2 and HSP60 phosphorylation in STn-expressing cells upon CD44s inhibition (**Fig. 7G**). STAT3 activation via CD44 has been linked to the acquisition of invasive traits and metastasis in different types of tumors (72, 73), suggesting a similar role in this context intimately linked to CD44s. Altogether, these observations provide glances on the intricate nature of oncogenic pathways regulation by CD44, which should be carefully addressed by dedicated phosphoproteomics studies in the future. Moreover, it portrayed the key role played by glycosylation in CD44s-mediated invasion of cancer cells. It suggests that these events may be modulated by sialylation of glycan chains, since this was not observed in Tn expressing cells. On the other hand, CD44s glycosylation appears to be key for defining the nature of oncogenic pathways supporting cancer aggressiveness.

In summary, we have highlighted that CD44s is a driver of invasion, as suggested by patient samples analysis showing a predominance of this isoform linked to more aggressive invasive tumors. Furthermore, we have shown that CD44s is implicated in an onset of relevant oncogenic pathways that support these functional traits. Finally, we found that the nature of the glycocode, most likely driven by sialylation, is crucial in this context and must be comprehensively investigated envisaging unequivocal identification by mass spectrometry, to fully disclose its functional and clinical relevance and provide means for precise cancer targeting.

## Discussion

CD44 is a pivotal glycoprotein in cancer, intricately related to most of its hallmarks, and widely accepted as a stem cell biomarker in BC (3, 16). The biological role of CD44 in cancer is intimately linked to the nature of protein isoforms (74) that, in the absence of dedicated analytical workflows, have been inferred from transcripts analysis complemented with immunoassays, lacking the necessary specificity for isoform distinction. Together with conflicting nomenclature, this has constituted a barrier for definitive CD44 exploitation in clinical settings. In fact, CD44 mRNA alternative splicing, which generates highly homologous polypeptide sequences and dense and heterogeneous glycosylation, has constituted a major obstacle for precise identification of cancer-specific glycoproteoforms and the establishment of solid structure-function relationships (2). Using BC as a model, we have started by exploring transcriptomics to provide the first glance at CD44 mosaicism in BC. This showed significant heterogeneity in terms of expressed isoforms and identified CD44s as a potential biomarker of invasion and poor prognosis. In agreement with these observations, CD44s was also strongly correlated with a subgroup of tumors displaying gene signatures characteristic of the basal BC molecular subtype but enriched for typical epithelial-to-mesenchymal transition markers. Later, we demonstrated that CD44s played an active role in supporting invasion *in vitro*, reinforcing the close link between this isoform and disease progression. Interestingly, basal bladder tumors express high levels of genes typical of more undifferentiated urothelial basal cells (*KRT5, KRT6*, and *KRT14)* together with several transcription factors that support stem cell homeostasis and cancer progression. The association of EMT markers with CSC properties has been reported for more aggressive tumors (75), which may likely be also the case for BC. In support of this hypothesis, in a recent study, Zhu *et al*. has elegantly demonstrated the pivotal role played by CD44s as regulator of stem-like properties, invasion, and lung metastasis in BC, providing solid molecular grounds to our current observations in patient samples (76) Similar reports have been found for breast (77), lung (78), hepatic (79), and pancreatic (11) cancers, which supports the pan-carcinogenic nature of CD44s and its role in regulating EMT and adaptive plasticity of CSC. Nevertheless, transcriptomics also showed that CD44s expression is not exclusive of tumors, as many healthy human organs may also express high levels of this isoform. This apparent lack of cancer specificity raises a major obstacle for precise cancer targeting, casting doubts on therapeutic development strategies. However, we also showed that targeting CD44s *O*-glycoforms, such as the Tn and STn antigens associated with aggressive forms of cancer, may allow to overcome this limitation.

Additionally, we tackled the limitations of transcriptomics and proteomics for definitive CD44 characterization, bringing together these different omics and glycomics. We have demonstrated that CD44 characterization cannot be achieved by a single omics and highlighted the importance of adding glycosylation for definitive isoforms identification, materializing the concept of glycoproteogenomics (28). Overall glycoproteogenomics analysis of different cell models and bladder tumors supported transcriptomics data and provided a mass spectrometry-based approach for validation of CD44 gene expression patterns. It also confirmed CD44s as a major carrier of altered glycosylation linked to cancer aggressiveness. Then, exploring the functional implications of CD44s in cancer cells, we concluded that this glycoprotein plays a pivotal role driving cell proliferation and invasion. Phosphoarray-based assays further reinforced the link between CD44s and key phospho-tyrosine kinases and related transcription factors, mainly associated to EGFR dependent signaling, supporting proliferation and invasion results and providing relevant molecular grounds to understand its role in BC progression. Similar observations have been made for other cancer models including gastric (80), breast (81), head and neck squamous cell carcinoma (54) and lung (55) tumors, reinforcing the influence of CD44 as a co-regulator of RTK signaling and EGFR mediated pathways, thereby contributing to cell proliferation and invasion. These findings support the relevance of addressing CD44 iso- and glycoforms in the context of precision oncology, namely for identification of patients benefiting from therapies targeting relevant oncogenic pathways.

Finally, exploring a library of glycoengineered cell models we have demonstrated that the nature of CD44s glycosylation is key to fine-tune its functional behavior, namely in terms of proliferation and invasion. CD44 is a heavily *O*-glycosylated protein, and it is likely that dramatic changes in glycans structures may induce major conformational changes implicating interactions with extracellular matrix components and cell surface receptors, which are yet to be fully understood (82). We found that CD44 inhibition in cells expressing high levels of sialylated *O*-glycans, including the STn antigen, had a major impact on cell invasion. Sialylation has long been described as a driver of cancer cells invasion, namely through the promotion of less cohesive cell phenotypes due to repulsive effects between proteins at the cell surface (83, 84). In fact, some reports suggest that sialylation could decrease the interaction of CD44 with its ligand hyaluronic acid, whose interaction contributes to cancer invasiveness and metastasis (25). By contrast, it has been previously demonstrated that CD44 *O*-glycans shortening increased hyaluronan binding capacity and may promote migratory pro-invasive cancer cell phenotypes in gastric cancer (82). Taken together, we propose that CD44s has a crucial role in cell invasion, however, it might be independent of the presence of STn moieties at its surface. On the other hand, a broader screening of main oncogenic pathways reflects another level of regulation dictated by the type of CD44s glycosylation, which appears to be key for defining oncogenic pathways adopted by cancer cells to support cancer aggressiveness. These findings, illustrate the relevance of CD44 glycoproteocode characterization towards true precision oncology and casts important research topics for addressing CD44 role in cancer. It also supports the implementation of systems biology approaches, namely through an interrogation of the phosphoproteome for more comprehensive understanding about the role of this glycoprotein in cancer.

In summary, we have provided molecular and functional contexts linking CD44s and, in particular, CD44 carrying immature *O*-glycosylation, to cancer aggressiveness, in agreement with its presence in invasive fronts and our previous reports linking both the protein and STn with invasion, metastasis and poor prognosis in BC (14, 36, 37). Therefore, we hypothesize that targeting CD44-Tn/STn glycoproteoforms may constitute a key strategy to control BC progression, which will be addressed in future studies. Given that CD44s and STn are frequently found in many human tumors linked to unfavorable prognosis (36, 85), we also believe that this may constitute a valuable cancer signature for future clinical interventions at different levels. Moreover, according to the analysis of healthy tissues, CD44s glycosignatures are highly cancer specific, holding tremendous potential to overcome some of the off-target limitations associated with glycans. Therefore, this study may come to contribute significantly to the development of antibodies and other ligands against cancer cells, enabling the creation of an onset of novel theranostics interventions (cancer detection, targeted therapeutics) and decisively aid patient stratification. For instance, encouraging pre-clinical studies were presented concerning CAR-Ts targeting Tn and STn antigens in MUC1 (86), which may now be adapted for more cancer specific proteoforms such as CD44-Tn/STn. Moreover, it may pave the way for glycan-based cancer vaccines capable of educating the immune system to respond to abnormally glycosylated cancer cells, avoiding cancer spread and relapse through protective memory. Overall, we have provided a roadmap to characterize the functional role of glycosylation in CD44, paving the way for more educated clinical interventions. This analytical strategy sets the grounds for a comprehensive characterization of many different types of tumors, constituting a crucial step towards understanding of the role of CD44 and other clinically relevant cell surface proteins in health and disease.

## Material and Methods

Details on materials and methods may be found in the SI Appendix Materials and Methods document. It includes information on BC patient samples, TCGA bioinformatics, cell lines and cell culture conditions, flow cytometry, splice variants analysis by NGS, CD44 immunoprecipitation, glycoproteogenomics protocol, nano-liquid chromatography-tandem mass spectrometry, bioinformatics analysis, western blot, real-time polymerase chain reaction, immunohistochemistry, immunofluorescence, proximity ligation assay, siRNA silencing assay, cell proliferation and invasion assays, phospho-kinase antibody array, and statistical analysis. Transcriptomics analysis of healthy tissues for CD44 isoforms were obtained from the GTEx Portal (https://gtexportal.org) on 03/06/21 and/or dbGaP accession number phs000424.v8.p2. The patient samples of bladder tumors were obtained from IPO-Porto biobank and used in this manuscript under the approval of the hospital’s ethics committee (project reference: CES 86/017) after obtaining informed patient’s consent.

## Data Availability

Proteomics datasets and tables describing protein assignment glycosylation annotations have been deposited in the Proteomics Identifications Database (PRIDE; https://www.ebi.ac.uk/pride/) and are freely available. All other study data are included in the article and/or SI Appendix.

## Acknowledgments

The authors wish to acknowledge the Portuguese Foundation for Science and Technology (FCT) for the human resources grants: PhD grants SFRH/BD/127327/2016 (CG), SFRH/BD/142479/2018 (JS), SFRH/BD/146500/2019 (MRS), SFRH/BD/111242/2015 (AP), 2020.08708.BD (RF), SFRH/BD/105355/2014 (RA), 2020.09384.BD (DF), and FCT assistant researcher grant CEECIND/03186/2017 (JAF). FCT is co-financed by European Social Fund (ESF) under Human Potential Operation Programme (POPH) from National Strategic Reference Framework (NSRF). The authors also acknowledge PhD Programs in Biomedical Sciences and Pathology and Molecular Genetics of ICBAS-University of Porto. The authors thank the financial support of the Portuguese Oncology Institute of Porto - Research Centre (CI-IPOP-29-2017-2020; CI-IPOP-58-2017-2021). This work was also financed by national funds through FCT/MCTES within the scope of the project UIDB/50006/2020 | UIDP/50006/2020, and by the Portuguese Mass Spectrometry Network, integrated in the National Roadmap of Research Infrastructures of Strategic Relevance (ROTEIRO/0028/2013; LISBOA-01-0145-FEDER-022125). The authors also wish to acknowledge the Genotype-Tissue Expression (GTEx) Project, supported by the Common Fund of the Office of the Director of the National Institutes of Health, and by NCI, NHGRI, NHLBI, NIDA, NIMH, and NINDS.

## Authors Contributions

CG, AMNS, JAF designed research; CG, JS, MRS, AP, DF, AB, EF, RA, PP, CP, LL, RF, AM, HO, AMNS performed research; CG, JS, MRS, AP, DF, AB, RA, LL, AMNS, LLS, JAF analyzed data; PP, HO, AMNS, LLS, JAF contributed new reagents/analytic tools; CG and JAF wrote the paper; all authors revised the paper.

