## Supporting Tables for "Glycoproteogenomics characterizes the CD44 splicing code driving bladder cancer invasion"

Cristiana Gaiteiro^a,b,c,d,e^, Janine Soares^a,b,c,d,f^, Marta Relvas-Santos^a,b,c,d,g,h,i^, Andreia Peixoto^a,b,c,g,h^, Dylan Ferreira^a,b,c,d,e,g,h,^, Andreia Brandão^b,c,j^, Elisabete Fernandes^a,b,c,k^, Rita Azevedo^l^, Paula Paulo^b,c,j^, Carlos Palmeira^a,b,c,k,m^, Luís Lima^a,b,c^, Rui Freitas^a,b,c,d,g,h^, Andreia Miranda^a,b,c,n^, Hugo Osório^g,n,o^, André M. N. Silva^i,p^, Jesús Prieto^e^, Lúcio Lara Santos^a,b,c,d,k,p,q^, José Alexandre Ferreira^a,b,c,d,p,1^

^a^Experimental Pathology and Therapeutics Group, IPO Porto Research Center (CI-IPOP), Portuguese Oncology Institute (IPO Porto), 4200-072 Porto, Portugal; ^b^RISE@CI-IPOP (Health Research Network), Portuguese Oncology Institute of Porto (IPO Porto), 4200-072 Porto, Portugal; ^c^Porto Comprehensive Cancer Center (P.ccc), 4200-072 Porto, Portugal; ^d^Institute of Biomedical Sciences Abel Salazar (ICBAS), University of Porto, 4050-013 Porto, Portugal; ^e^Center for Applied Medical Research (Centro de Investigación Médica Aplicada, CIMA), University of Navarra, 31008 Pamplona, Navarra, Spain; ^f^REQUIMTE-LAQV, Department of Chemistry, University of Aveiro, 3810-193 Aveiro, Portugal; ^g^Institute for Research and Innovation in Health (i3S), University of Porto, 4200-135 Porto, Portugal; ^h^Institute for Biomedical Engineering (INEB), University of Porto, 4200-135 Porto, Portugal; ^i^REQUIMTE-LAQV, Department of Chemistry and Biochemistry, Faculty of Sciences of the University of Porto, 4169-007 Porto, Portugal; ^j^Cancer Genetics Group, IPO Porto Research Center (CI-IPOP), Portuguese Oncology Institute (IPO Porto), 4200-072 Porto, Portugal; ^k^FP-I3ID, University Fernando Pessoa, 4249-004 Porto, Portugal; ^l^Laboratoire d'Etude du Métabolisme des Médicaments (LEMM), CEA, INRA, Université Paris Saclay, F-91191, Gif-sur-Yvette cedex, France.; ^m^Immunology Department, Portuguese Oncology Institute (IPO Porto), 4200-072 Porto, Portugal; ^n^Faculty of Medicine of the University of Porto, 4200-319 Porto, Portugal; ^o^Ipatimup—Institute of Molecular Pathology and Immunology of the University of Porto, University of Porto, 4200-135 Porto, Portugal; ^p^GlycoMatters Biotech, 4500-162 Espinho, Portugal; ^q^Department of Surgical Oncology, Portuguese Oncology Institute (IPO Porto), 4200-072 Porto, Portugal

**^1^Corresponding author:**

José Alexandre Ferreira

ORCID ID: https://orcid.org/0000-0002-0097-6148

Experimental Pathology and Therapeutics Group, Research Centre, Portuguese Oncology Institute of Porto, R. Dr. António Bernardino de Almeida 62, 4200-072 Porto, Portugal; Tel. +351 225084000 (ext. 5111).

**Running head:** CD44 glycoforms in bladder cancer

**Keywords:** glycomics; proteogenomics; glycoproteogenomics; bladder cancer; CD44

**Table S1. Tailored-made Taqman Gene Expression Assay ID’s to specifically detect mRNA encoding for total CD44 and its 4 splicing variants.**

| Gene | TaqMan Gene Expression Assay ID |
| --- | --- |
| CD44 Total | Hs01075864_m1 |
| CD44v2-10 | Hs01075866_m1 |
| CD44v3-10 | Hs01081480_m1 |
| CD44v8-10 | Hs01081475_m1 |
| CD44s/st | Hs01081473_m1 |

**Table S2. CD44 isoforms identified in 5637 and T24 cells by whole transcriptome analysis.**

| Ensembl | Uniprot | CD44 Isoforms |
| --- | --- | --- |
| ENST00000263398.10 | P16070-12 | **CD44s** |
| [ENST00000526025.2](https://www.ensembl.org/Homo_sapiens/Transcript/Summary?db=core;g=ENSG00000026508;r=11:35138882-35232402;t=ENST00000526025) | P16070-2 or E9PKC6 | **CD44spliceE2** |
| ENST00000415148.6 | P16070-3 | **CD44v3-10** |
| ENST00000428726.6 | P16070-1 | **CD44v2-10** |
| ENST00000433892.6 | P16070-10 | **CD44v8-10** |
| ENST00000278386.10 | P16070-19 | **CD44sol** |
| ENST00000434472.6 | P16070-11 | **CD44v10** |
| ENST00000352818.8 | P16070-18 | **CD44s-exon15** |
| ENST00000442151.6 | P16070-15 or H0Y5E4 | **CD44st** |
| [ENST00000526669.6](https://www.ensembl.org/Homo_sapiens/Transcript/Summary?db=core;g=ENSG00000026508;r=11:35138882-35232402;t=ENST00000526669) | H0YD13 | - |
| [ENST00000425428.6](https://www.ensembl.org/Homo_sapiens/Transcript/Summary?db=core;g=ENSG00000026508;r=11:35138882-35232402;t=ENST00000425428) | Q86UZ1 | - |
| [ENST00000528086.5](https://www.ensembl.org/Homo_sapiens/Transcript/Summary?db=core;g=ENSG00000026508;r=11:35138882-35232402;t=ENST00000528086) | - | - |
| [ENST00000526000.6](https://www.ensembl.org/Homo_sapiens/Transcript/Summary?db=core;g=ENSG00000026508;r=11:35138882-35232402;t=ENST00000526000) | H0YDW7 | - |
| [ENST00000279452.10](https://www.ensembl.org/Homo_sapiens/Transcript/Summary?db=core;g=ENSG00000026508;r=11:35138882-35232402;t=ENST00000279452) | H0Y2P0 | - |
| [ENST00000531118.5](https://www.ensembl.org/Homo_sapiens/Transcript/Summary?db=core;g=ENSG00000026508;r=11:35138882-35232402;t=ENST00000531118) | - | - |
| [ENST00000528455.5](https://www.ensembl.org/Homo_sapiens/Transcript/Summary?db=core;g=ENSG00000026508;r=11:35138882-35232402;t=ENST00000528455) | H0YD17 | - |
| [ENST00000531873.5](https://www.ensembl.org/Homo_sapiens/Transcript/Summary?db=core;g=ENSG00000026508;r=11:35138882-35232402;t=ENST00000531873) | H0YD90 | - |
| [ENST00000525209.5](https://www.ensembl.org/Homo_sapiens/Transcript/Summary?db=core;g=ENSG00000026508;r=11:35138882-35232402;t=ENST00000525209) | - | - |

**Table S3. Glycopeptides identified by nanoLC-HCD/CID-MS/MS in glycoproteogenomics settings for 5637 cells.**

**Table S4. Glycopeptides identified by nanoLC-HCD/CID-MS/MS in glycoproteogenomics settings for T24 cells.**

**Table S5. Glycopeptides identified by nanoLC-HCD/CID-MS/MS in glycoproteogenomics settings for CD44s^high^ MIBC showing areas of CD44 and STn co-localization.**

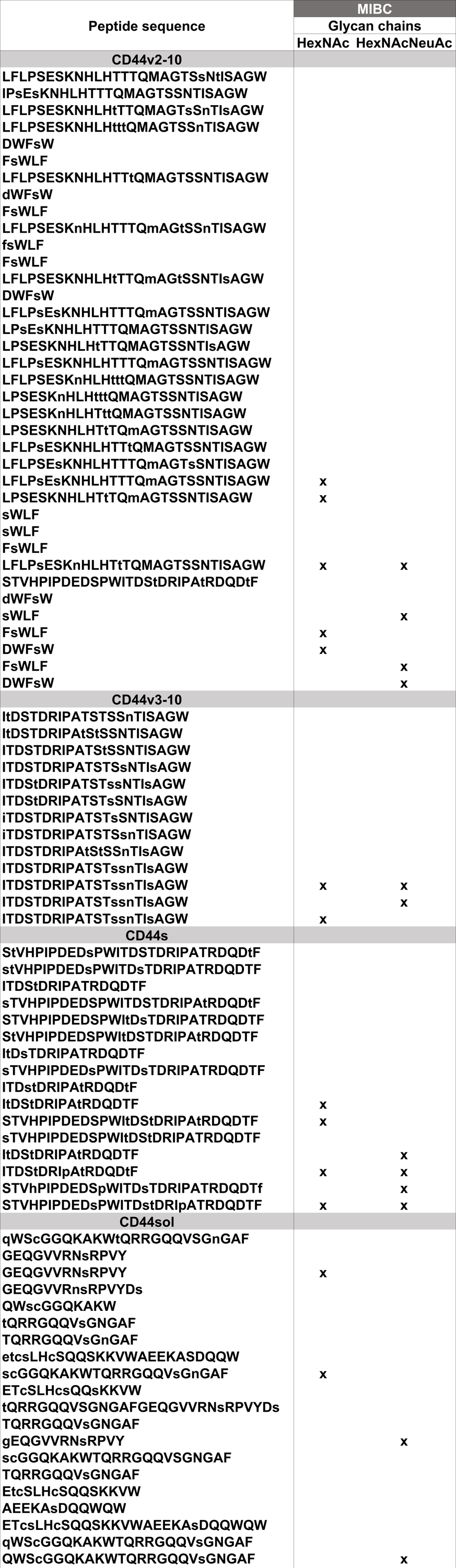

**Table S6. Glycopeptides identified by nanoLC-HCD/CID-MS/MS in glycoproteogenomics settings for T24 *C1GALT1* KO cell model**

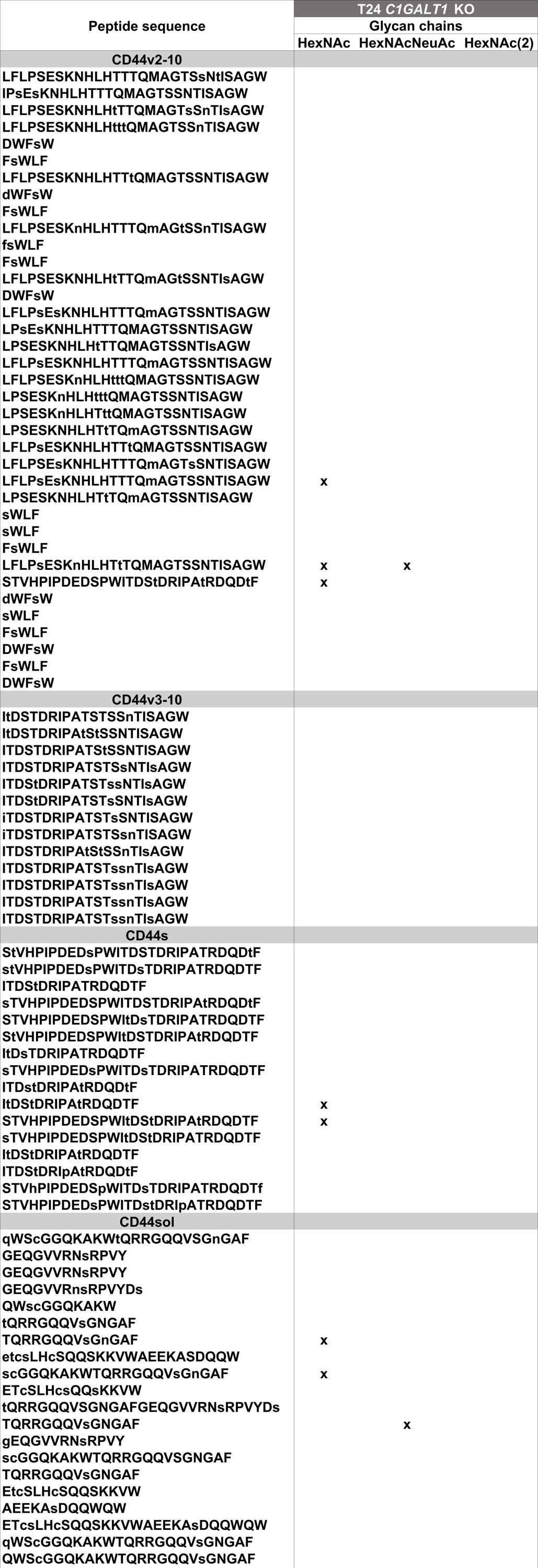
