## Supporting Figures for "Glycoproteogenomics characterizes the CD44 splicing code driving bladder cancer invasion"

Cristiana Gaiteiro^a,b,c,d,e^, Janine Soares^a,b,c,d,f^, Marta Relvas-Santos^a,b,c,d,g,h,i^, Andreia Peixoto^a,b,c,g,h^, Dylan Ferreira^a,b,c,d,e,g,h,^, Andreia Brandão^b,c,j^, Elisabete Fernandes^a,b,c,k^, Rita Azevedo^l^, Paula Paulo^b,c,j^, Carlos Palmeira^a,b,c,k,m^, Luís Lima^a,b,c^, Rui Freitas^a,b,c,d,g,h^, Andreia Miranda^a,b,c,n^, Hugo Osório^g,n,o^, André M. N. Silva^i,p^, Jesús Prieto^e^, Lúcio Lara Santos^a,b,c,d,k,p,q^, José Alexandre Ferreira^a,b,c,d,p,1^

^a^Experimental Pathology and Therapeutics Group, IPO Porto Research Center (CI-IPOP), Portuguese Oncology Institute (IPO Porto), 4200-072 Porto, Portugal; ^b^RISE@CI-IPOP (Health Research Network), Portuguese Oncology Institute of Porto (IPO Porto), 4200-072 Porto, Portugal; ^c^Porto Comprehensive Cancer Center (P.ccc), 4200-072 Porto, Portugal; ^d^Institute of Biomedical Sciences Abel Salazar (ICBAS), University of Porto, 4050-013 Porto, Portugal; ^e^Center for Applied Medical Research (Centro de Investigación Médica Aplicada, CIMA), University of Navarra, 31008 Pamplona, Navarra, Spain; ^f^REQUIMTE-LAQV, Department of Chemistry, University of Aveiro, 3810-193 Aveiro, Portugal; ^g^Institute for Research and Innovation in Health (i3S), University of Porto, 4200-135 Porto, Portugal; ^h^Institute for Biomedical Engineering (INEB), University of Porto, 4200-135 Porto, Portugal; ^i^REQUIMTE-LAQV, Department of Chemistry and Biochemistry, Faculty of Sciences of the University of Porto, 4169-007 Porto, Portugal; ^j^Cancer Genetics Group, IPO Porto Research Center (CI-IPOP), Portuguese Oncology Institute (IPO Porto), 4200-072 Porto, Portugal; ^k^FP-I3ID, University Fernando Pessoa, 4249-004 Porto, Portugal; ^l^Laboratoire d'Etude du Métabolisme des Médicaments (LEMM), CEA, INRA, Université Paris Saclay, F-91191, Gif-sur-Yvette cedex, France.; ^m^Immunology Department, Portuguese Oncology Institute (IPO Porto), 4200-072 Porto, Portugal; ^n^Faculty of Medicine of the University of Porto, 4200-319 Porto, Portugal; ^o^Ipatimup—Institute of Molecular Pathology and Immunology of the University of Porto, University of Porto, 4200-135 Porto, Portugal; ^p^GlycoMatters Biotech, 4500-162 Espinho, Portugal; ^q^Department of Surgical Oncology, Portuguese Oncology Institute (IPO Porto), 4200-072 Porto, Portugal

**^1^Corresponding author:**

José Alexandre Ferreira

ORCID ID: https://orcid.org/0000-0002-0097-6148

Experimental Pathology and Therapeutics Group, Research Centre, Portuguese Oncology Institute of Porto, R. Dr. António Bernardino de Almeida 62, 4200-072 Porto, Portugal; Tel. +351 225084000 (ext. 5111).

**Running head:** CD44 glycoforms in bladder cancer

**Keywords:** glycomics; proteogenomics; glycoproteogenomics; bladder cancer; CD44

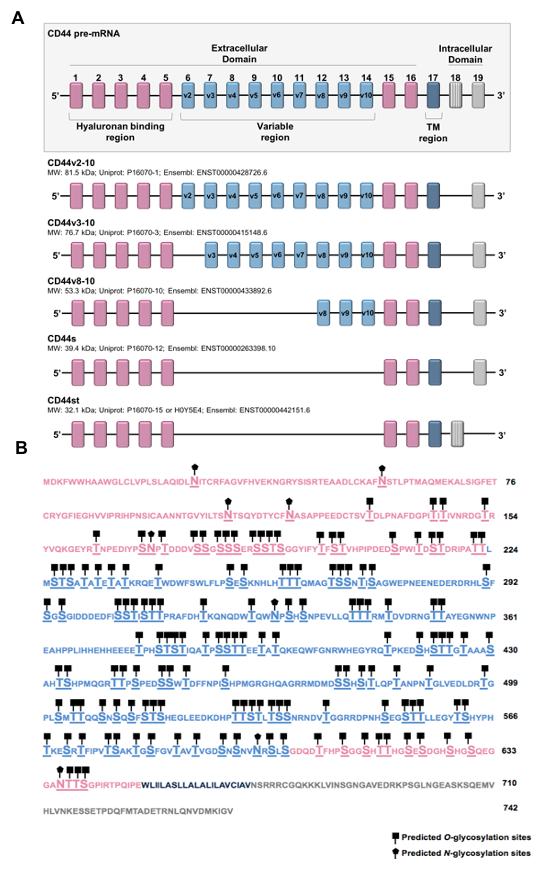

**Fig. S1. A) Schematic representation of human CD44 pre-mRNA and experimentally confirmed isoforms resulting from alternative splicing and extensively explored in this study.** Exons generally regarded as conserved are represented in pink (exons 1-5 and 15-16), variable exons are represented in blue (exons 6-14), the transmembrane region is represented in dark blue (exon 17), and the intracellular tail is represented in grey (exons 18-19). Exon 18, filled with gray stripes, contains an early 3’UTR and is only present in the CD44st isoform. The estimated molecular weight for each variant (without posttranslational modifications), Uniprot and Ensemble codifications are also highlighted. **B) Amino acid sequence evidencing potential *O*- and *N*-glycosites.** *O*-glycosylation sites were predicted using the Net*O*Glyc 4.0 server (http://www.cbs.dtu.dk/services/NetOGlyc/) and *N-glycosylation sites with the NetNGlyc 1.0 server* (http://www.cbs.dtu.dk/services/NetNGlyc/). The different domains of the protein are highlighted using the same color code as in panel A and evidences the high density of *O*-glycosites in comparison to *N*-glycosites, which is particularly evident in the variable region.

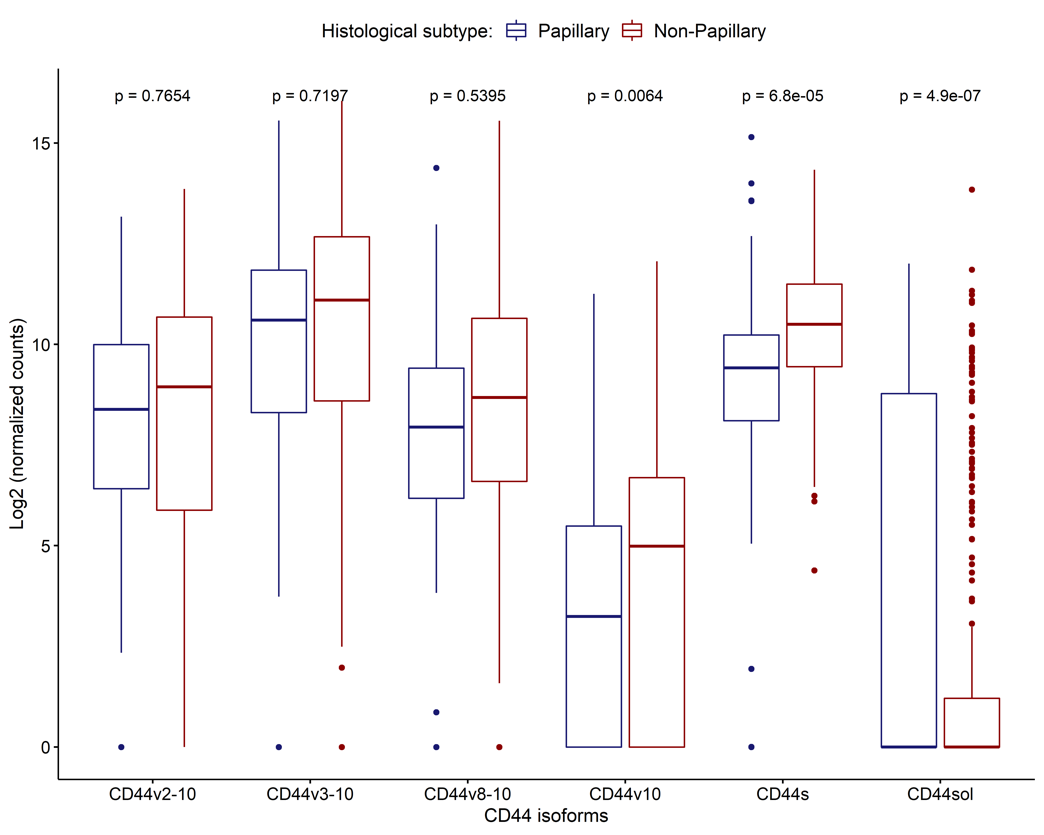

**Fig. S2. CD44 variants expression according to the histological subtypes of TCGA muscle invasive bladder tumors.** Wilcoxon test revealed a significant overexpression of CD44s in more aggressive non-papillary lesions.

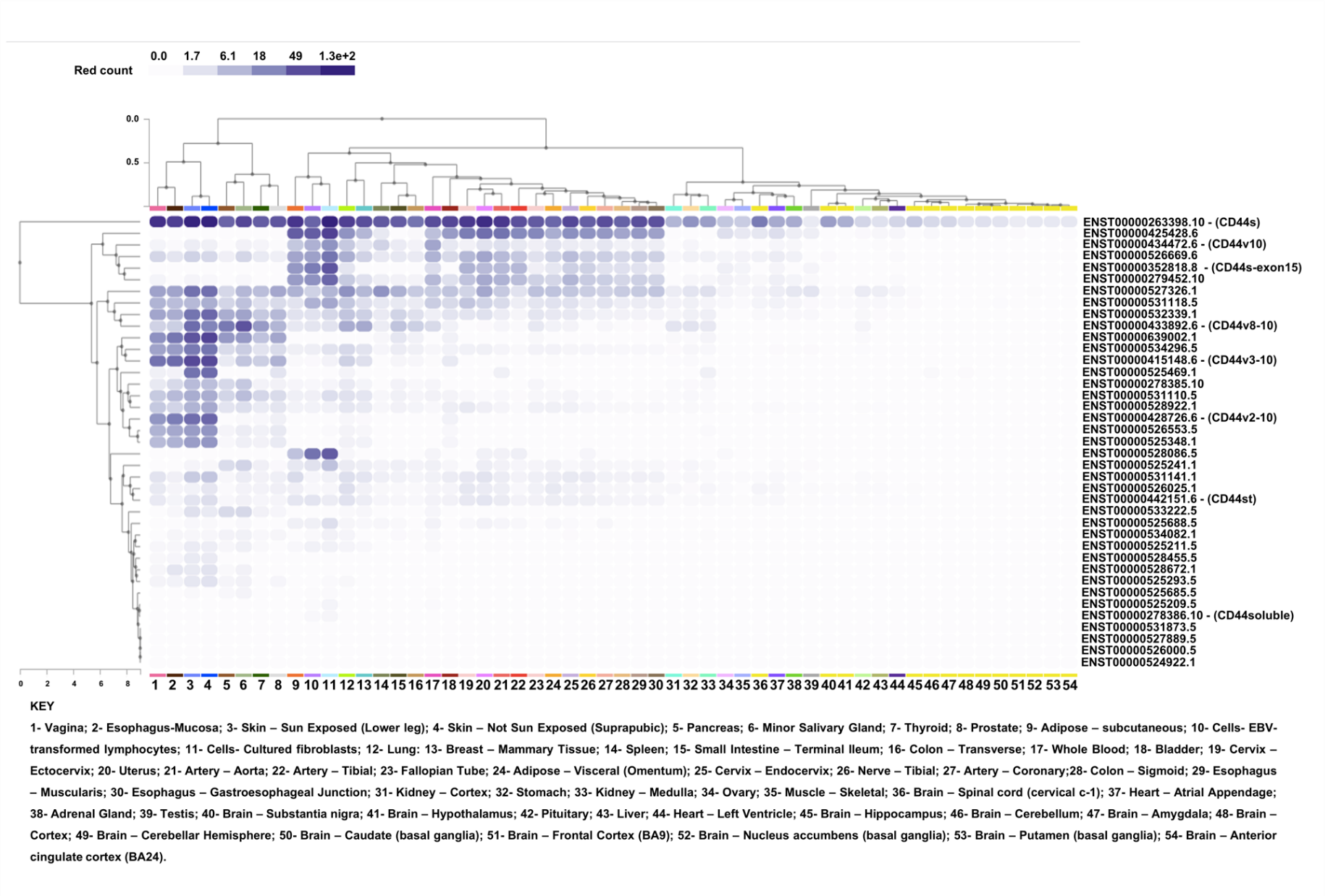

**Fig. S3. CD44 splice variants expression in healthy human tissues.** Transcriptomics analysis of healthy tissues for CD44 isoforms were obtained and adapted from the GTEx Portal on 03/26/21 and/or dbGaP accession number phs000424.v8.p2. The Figure shows the complex mosaicism of CD44 mRNA in healthy tissues, including the co-expression of multiple isoforms in the same tissue. It also highlights that most healthy tissues/non-neoplastic cells express high levels of CD44s.

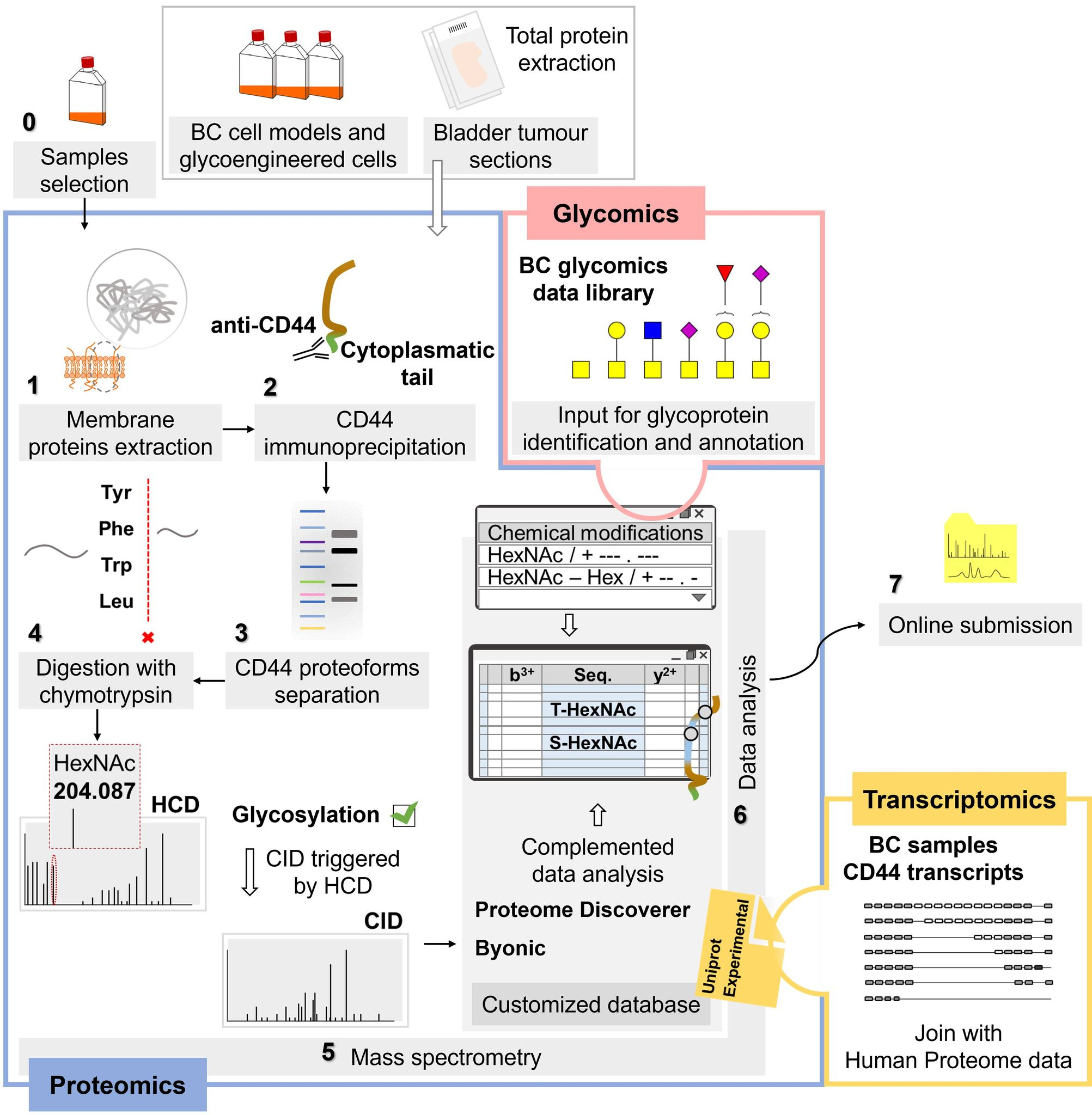

**Fig. S4. Glycoproteogenomics workflow for identification of CD44 glycoproteoforms.** Briefly, bladder cancer cell models, glycoengineered cells and tumor tissue sections were used as starting material. **1.** Membrane glycoproteins from cell lines are isolated by differential ultracentrifugation. **2.** CD44 is immunoprecipitated using a monoclonal antibody targeting the cytoplasmatic tail domain expressed by most CD44 isoforms. **3.** CD44-IP enriched extracts were then digested with sialidase, separated by gradient SDS-PAGE electrophoresis. **4.** Bands are excised from the gels, proteins are reduced, alkylated, and digested with chymotrypsin *in* gel. **5.** Chymotrypsin digests are analyzed by nanoLC-MS/MS (HCD-MS2 and CID-MS2 with a dependent acquisition based on HCD detection of HexNAc oxonium ion at *m/z* 204.087. **6.** Glycoproteoforms identification using a glycoproteogenomics approach. RNAseq was used to identify CD44 isoforms expressed by 5637 and T24 cells. Predicted protein sequences were included in a database together with the human proteome from Uniprot database and used for protein identification in the Proteome Discoverer bioinformatics platform. Glycomics analysis was performed in parallel to determine the glycan structures that may be present as variable CD44 post-translational modification. This information was included upon protein identification and glycopeptides annotation by MS/MS.

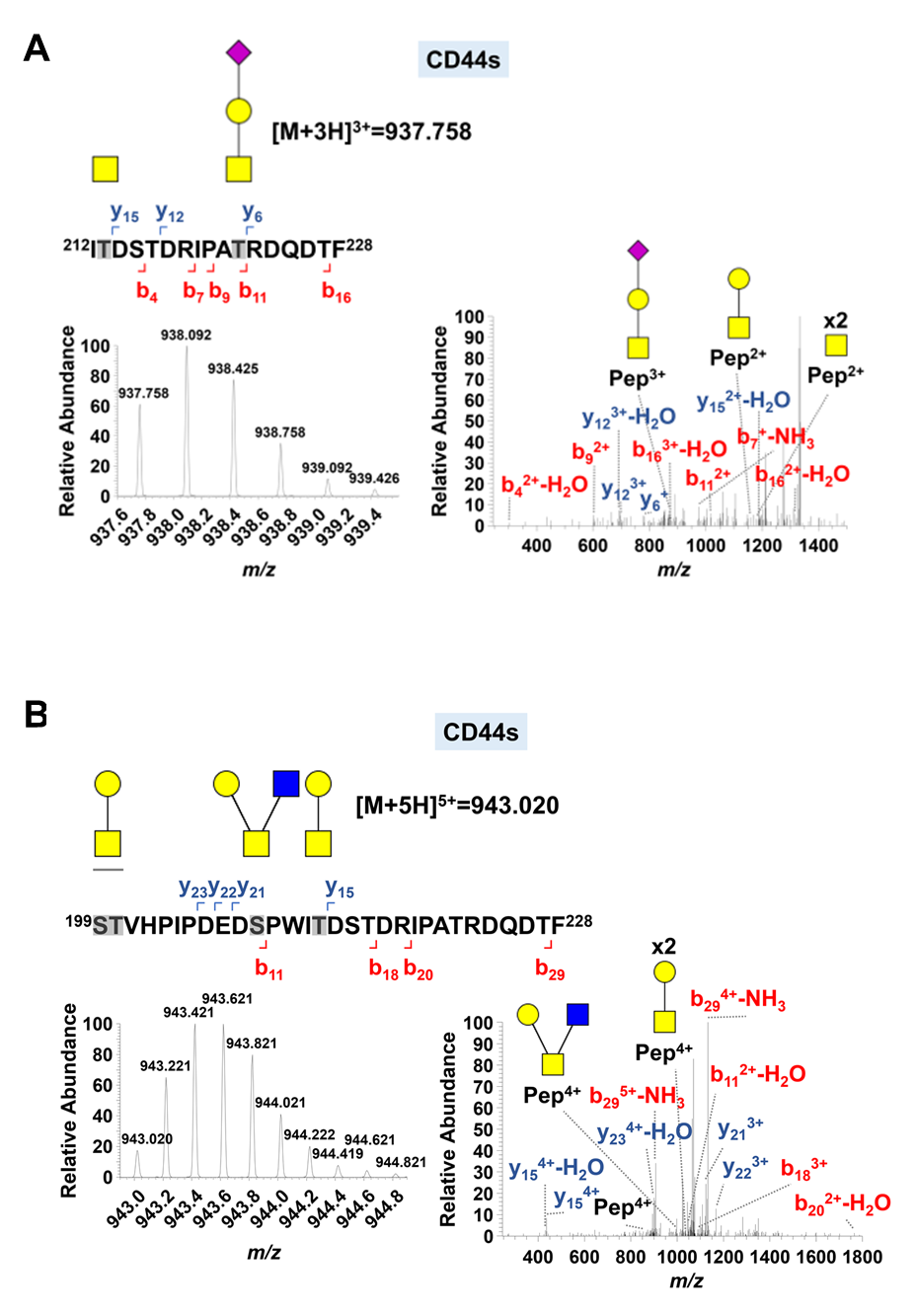

**Fig. S5.** **CID spectra for CD44s glycopeptides carrying *O*-glycans identified in T24 cells.** **A) CID spectrum for CD44s-Tn/ST and B) CD44s-T/core 2 glycopeptides.** CD44s glycopeptides carrying these *O*-glycans were identified by CID based on an HCD data dependent acquisition. Briefly, species presenting HCD product ion spectrum with an HexNAc fragment at m/z 204.087 were elected for CID fragmentation. CID spectra confirmed the glycan structures and presented several *y-* and *b-*series peptide fragments that helped supporting glycosites assignment (highlighted in grey in the peptide sequence). Collectively, these spectra confirm the presence of the illustrated *O*-glycans in CD44s from T24 cells and the heterogeneous nature of glycosylation at the peptide level.

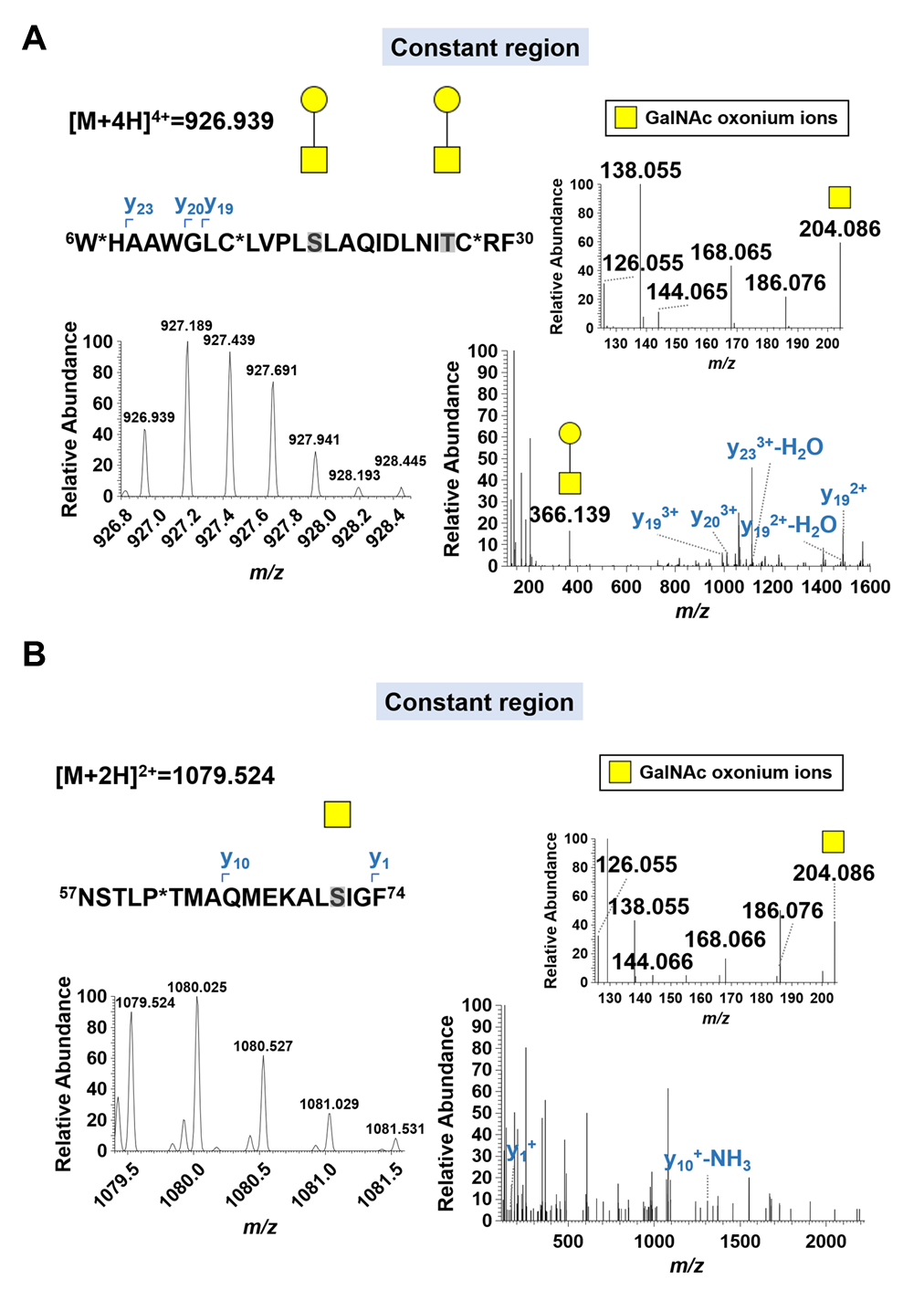

**Fig. S6.** **A) CD44-T and B) Tn glycopeptides belonging to the conserved extracellular domain in different isoforms identified in muscle invasive bladder tumors by nanoLC-HCD-MS/MS.** Both MS/MS spectra present the HexNAc oxonium ions, namely the ion at *m/z* 204.086. The MS/MS spectrum of the CD44-T glycopeptide also presents an evident ion at *m/z* 366.139, corresponding to the oxonium ion for the T antigen. y-series ions resulting from peptide backbone fragmentation were also highlighted. The sequence of the glycopeptide presents the glycosites identified in grey. The symbol *corresponds to amino acid modifications: W – hydroxykynurenine; C – carbamidomethyl; P – glutamic semialdehyde.

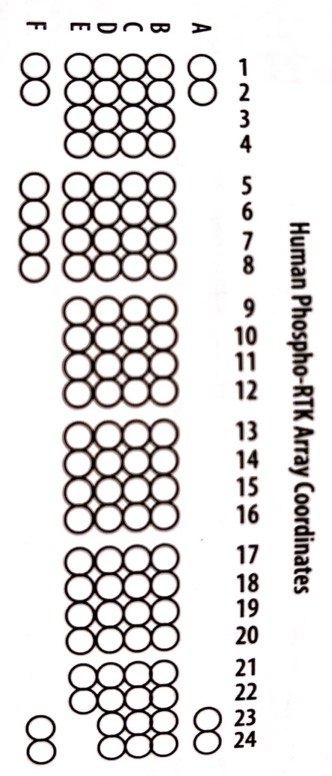

| Coordinate | Receptor Family | RTK/Control |
| --- | --- | --- |
| A1, A2 | Reference Spots | - |
| B1, B2 | Reference Spots | - |
| B3, B4 | EGF R | EGF R |
| B5, B6 | EGF R | ErbB2 |
| B7, B8 | EGF R | ErbB4 |
| B9, B10 | FGF R | FGF R1 |
| B11, B12 | FGF R | FGF R2α |
| B13,B14 | FGF R | FGF R3 |
| B15, B16 | FGF R | FGF R4 |
| B17, B18 | Insulin R | Insulin R |
| B19, B20 | Insulin R | IGF-I R |
| B21, B22 | Axl | Axl |
| B23, B24 | Axl | Dtk |
| C1, C2 | Axl | Mer |
| C3, C4 | HGF R | HGF R |
| C5, C6 | HGF R | MSP R |
| C7, C8 | PDGF R | PDGF Rα |
| C9, C10 | PDGF R | PDGF Rβ |
| C11, C12 | PDGF R | SCF R |
| C13, C14 | PDGF R | Flt-3 |
| C15, C16 | PDGF R | M-CSF R |
| C17, C18 | RET | c-Ret |
| C19, C20 | ROR | ROR1 |
| C21, C22 | ROR | ROR2 |
| C23, C24 | Tie | Tie-1 |
| D1, D2 | Tie | Tie-2 |
| D3, D4 | NGF R | TrkA |
| D5, D6 | NGF R | TrkB |
| D7, D8 | NGF R | TrkC |
| D9, D10 | VEGF R | VEGF R1 |
| D11, D12 | VEGF R | VEGF R2 |
| D13, D14 | VEGF R | VEGF R3 |
| D15, D16 | MuSK | MuSK |
| D17, D18 | Eph R | EphA1 |
| D19, D20 | Eph R | EphA2 |
| D21, D22 | Eph R | EphA3 |
| D23, D24 | Eph R | EphA4 |
| E1, E2 | Eph R | EphA6 |
| E3, E4 | Eph R | EphA7 |
| E5, E6 | Eph R | EphB1 |
| E7, E8 | Eph R | EphB2 |
| E9, E10 | Eph R | EphB4 |
| E11, E12 | Eph R | EphB6 |
| E13, E14 | Insulin R | ALK |
| E15, E16 | - | DDR1 |
| E17, E18 | - | DDR2 |
| E19, E20 | Eph R | EphA5 |
| E21, E22 | Eph R | EphA10 |
| F1, F2 | Reference spots | - |
| F5, F6 | Eph R | Ephb3 |
| F7, F8 | - | RYK |
| F23, F24 | Control (-) | PBS |

**Fig. S7. The human phospho-receptor tyrosine kinase (Phospho-RTK) array coordinates.** The phospho-array presents capture and control antibodies spotted in duplicates for the 49 different phosphorylated human RTKs, which are represented by alpha-numeric coordinates.

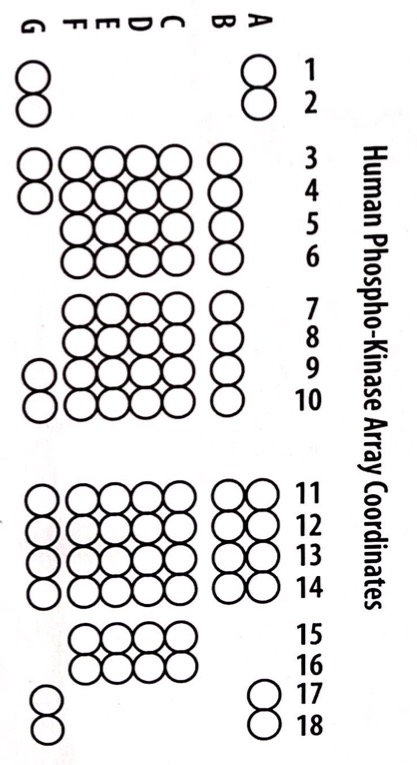

| Coordinate | Target/Control | Phosphorylation Site |
| --- | --- | --- |
| A1, A2 | Reference Spot | - |
| A11,A12 | Akt 1/2/3 | T308 |
| A13, A14 | Akt 1/2/3 | S473 |
| A17, A18 | Reference Spot | - |
| B3, B4 | CREB | S133 |
| B5, B6 | EGF R | Y1086 |
| B7, B8 | eNOS | S1177 |
| B9, B10 | ERK 1/2 | T202/Y204, T185/Y187 |
| B11, B12 | Chk-2 | T68 |
| B13, B14 | c-Jun | S63 |
| C3, C4 | Fgr | Y412 |
| C5, C6 | GSK-3α/β | S21/S9 |
| C7, C8 | GSK-3β | S9 |
| C9, C10 | HSP27 | S78/S82 |
| C11, C12 | p53 | S15 |
| C13, C14 | p53 | S46 |
| C15, C16 | p53 | S392 |
| D3, D4 | JNK 1/2/3 | T183/Y185, T221/Y223 |
| D5, D6 | Lck | Y394 |
| D7, D8 | Lyn | Y397 |
| D9, D10 | MSK 1/2 | S376/S360 |
| D11, D12 | p70 S6 Kinase | T389 |
| D13, D14 | p70 S6 Kinase | T421/S424 |
| D15, D16 | PRAS40 | T246 |
| E3, E4 | p38α | T180/Y182 |
| E5, E6 | PDGF Rβ | Y751 |
| E7, E8 | PLC-γ1 | Y783 |
| E9, E10 | Src | Y419 |
| E11, E12 | PYK2 | Y402 |
| E13, E14 | RSK 1/2 | S221/S227 |
| E15, E16 | RSK 1/2/3 | S380/S386/S377 |
| F3, F4 | STAT2 | Y689 |
| F5, F6 | STAT5a/b | Y694/Y699 |
| F7, F8 | WNK1 | T60 |
| F9, F10 | Yes | Y426 |
| F11, F12 | STAT1 | Y701 |
| F13, F14 | STAT3 | Y705 |
| F15, F16 | STAT3 | S727 |
| G1, G2 | Reference Spot | - |
| G3, G4 | β-Catenin | - |
| G9, G10 | PBS (Control -) | - |
| G11, G12 | STAT6 | Y641 |
| G13, G14 | HSP60 | - |
| G17, G18 | PBS (Control -) | - |

**Fig. S8. The human phospho-kinase array coordinates.** The phospho-array presents capture and control antibodies spotted in duplicates for the detection of the relative levels of phosphorylation of 37 kinase phosphorylation sites and 2 related total proteins represented by alpha-numeric coordinates.

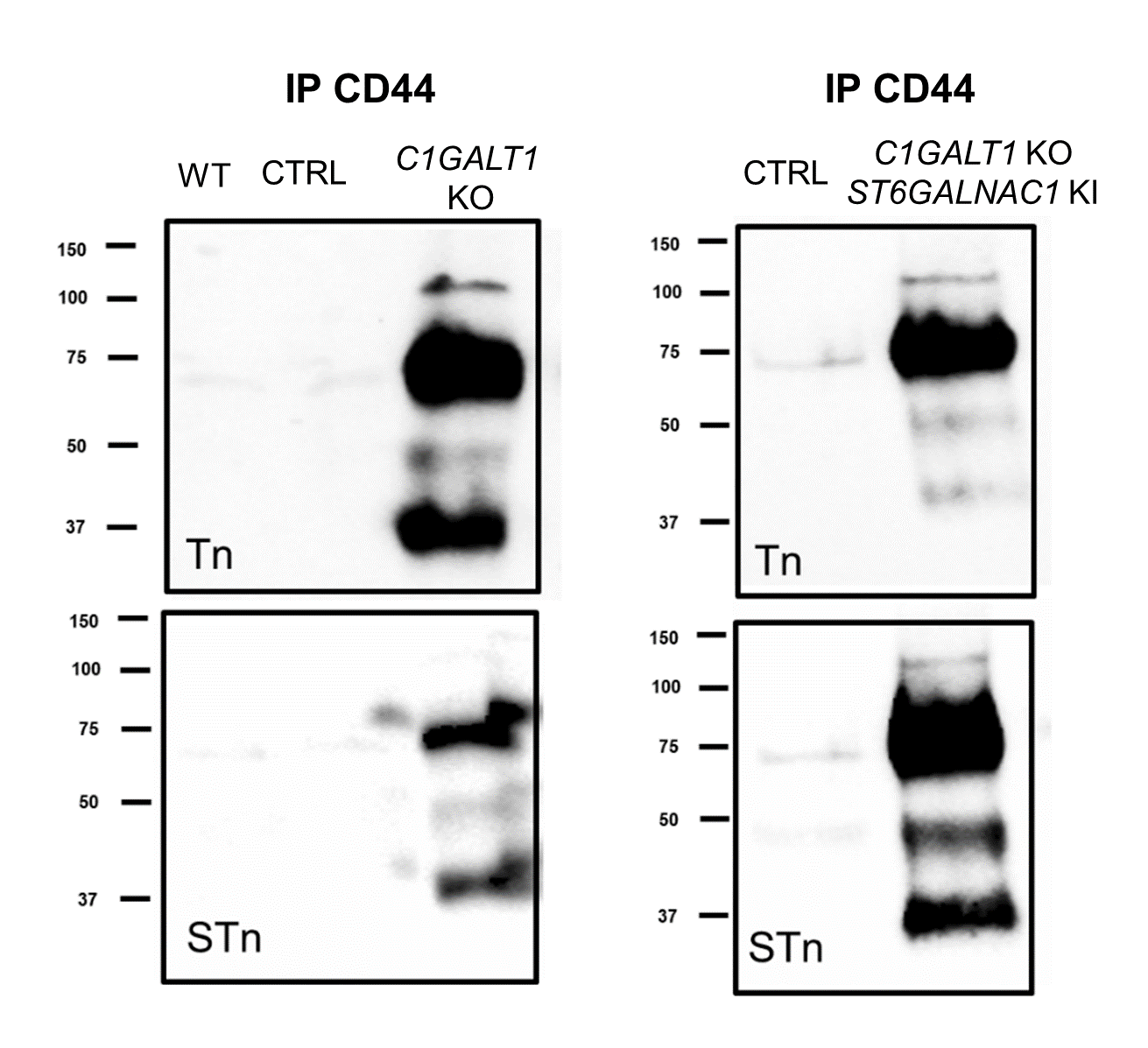

**Fig. S9. Western Blots show co-expression of CD44 and short-chain *O*-glycans, namely STn and Tn, in glycoengineered T24 cells.** CD44 was immunoprecipitated from membrane protein extracts of T24 *C1GALT1*KO, T24 *C1GALT1*KO/*ST6GALNAC1*KI and corresponding controls and then blotted for STn, Tn, and CD44. A demarked expression of CD44-STn and CD44-Tn was detected for both cell model mainly in bands at 75 kDa and bellow.

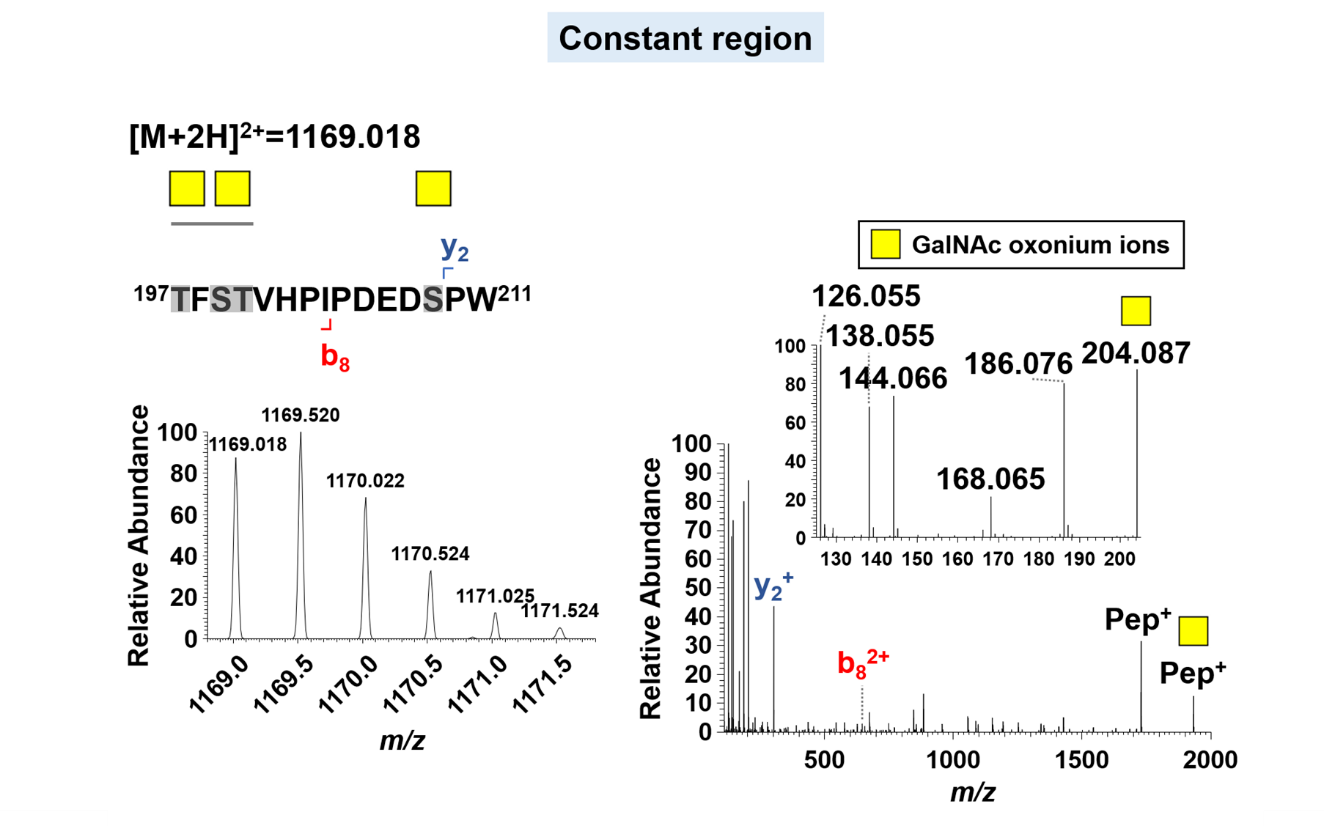

**Fig. S10.** **CD44-Tn glycopeptides belonging to the conserved extracellular domain identified in T24 *C1GALT1*KO cells by nanoLC-HCD-MS/MS.** MS/MS spectrum highlights the typical HexNAC oxonium ion (*m/z* 204.087) as well as other oxonium ion fragments consistent with GalNAc (highlighted at *m/z* 138.055 and 144.066). The MS/MS spectrum of the CD44-Tn glycopeptide also highlights peptide ions with GalNAc losses. The predicted and possible glycosites are identified in grey.
