## Supplementary Materials for "Glycoproteogenomics characterizes the CD44 splicing code driving bladder cancer invasion"

**-Supplementary Material and Methods-**

Cristiana Gaiteiro^a,b,c,d,e^, Janine Soares^a,b,c,d,f^, Marta Relvas-Santos^a,b,c,d,g,h,i^, Andreia Peixoto^a,b,c,g,h^, Dylan Ferreira^a,b,c,d,e,g,h,^, Andreia Brandão^b,c,j^, Elisabete Fernandes^a,b,c,k^, Rita Azevedo^l^, Paula Paulo^b,c,j^, Carlos Palmeira^a,b,c,k,m^, Luís Lima^a,b,c^, Rui Freitas^a,b,c,d,g,h^, Andreia Miranda^a,b,c,n^, Hugo Osório^g,n,o^, André M. N. Silva^i,p^, Jesús Prieto^e^, Lúcio Lara Santos^a,b,c,d,k,p,q^, José Alexandre Ferreira^a,b,c,d,p,1^

^a^Experimental Pathology and Therapeutics Group, IPO Porto Research Center (CI-IPOP), Portuguese Oncology Institute (IPO Porto), 4200-072 Porto, Portugal; ^b^RISE@CI-IPOP (Health Research Network), Portuguese Oncology Institute of Porto (IPO Porto), 4200-072 Porto, Portugal; ^c^Porto Comprehensive Cancer Center (P.ccc), 4200-072 Porto, Portugal; ^d^Institute of Biomedical Sciences Abel Salazar (ICBAS), University of Porto, 4050-013 Porto, Portugal; ^e^Center for Applied Medical Research (Centro de Investigación Médica Aplicada, CIMA), University of Navarra, 31008 Pamplona, Navarra, Spain; ^f^REQUIMTE-LAQV, Department of Chemistry, University of Aveiro, 3810-193 Aveiro, Portugal; ^g^Institute for Research and Innovation in Health (i3S), University of Porto, 4200-135 Porto, Portugal; ^h^Institute for Biomedical Engineering (INEB), University of Porto, 4200-135 Porto, Portugal; ^i^REQUIMTE-LAQV, Department of Chemistry and Biochemistry, Faculty of Sciences of the University of Porto, 4169-007 Porto, Portugal; ^j^Cancer Genetics Group, IPO Porto Research Center (CI-IPOP), Portuguese Oncology Institute (IPO Porto), 4200-072 Porto, Portugal; ^k^FP-I3ID, University Fernando Pessoa, 4249-004 Porto, Portugal; ^l^Laboratoire d'Etude du Métabolisme des Médicaments (LEMM), CEA, INRA, Université Paris Saclay, F-91191, Gif-sur-Yvette cedex, France.; ^m^Immunology Department, Portuguese Oncology Institute (IPO Porto), 4200-072 Porto, Portugal; ^n^Faculty of Medicine of the University of Porto, 4200-319 Porto, Portugal; ^o^Ipatimup—Institute of Molecular Pathology and Immunology of the University of Porto, University of Porto, 4200-135 Porto, Portugal; ^p^GlycoMatters Biotech, 4500-162 Espinho, Portugal; ^q^Department of Surgical Oncology, Portuguese Oncology Institute (IPO Porto), 4200-072 Porto, Portugal

**^1^Corresponding author:**

José Alexandre Ferreira

ORCID ID: https://orcid.org/0000-0002-0097-6148

Experimental Pathology and Therapeutics Group, Research Centre, Portuguese Oncology Institute of Porto, R. Dr. António Bernardino de Almeida 62, 4200-072 Porto, Portugal; Tel. +351 225084000 (ext. 5111).

**Running head:** CD44 glycoforms in bladder cancer

**Keywords:** glycomics; proteogenomics; glycoproteogenomics; bladder cancer; CD44

**Material and Methods**

**Patients sampling and healthy human tissues**

A retrospective series of 75 formalin-fixed paraffin-embedded (FFPE) bladder tumor tissues from the Portuguese Institute of Oncology of Porto (IPO-Porto) biobank were used for this study. Bladder tumors were surgically removed from 61 male and 14 female, ranging from 26 to 85 years of age (67 ± 11.7), admitted and treated at the IPO-Porto between 2000 and 2017. The tumors were classified as non-muscle invasive (≤T1, NMIBC; n=34) and muscle-invasive (≥T2; MIBC; n=41) bladder cancers. Eleven histologically normal urothelium tissue sections from healthy individuals were included. Additionally, a broad library of healthy tissues (liver, colon, small intestine, gallbladder, pancreas, thyroid, stomach, appendix, testicle, skin, breast, kidney, lung, mucosa-associated lymphoid tissue-MALT, and white blood cells) were also considered. Bladder tumor and urothelial sections were characterized in terms of CD44 isoforms by RT-PCR and glycoproteogenomics. Additionally, tumors and healthy tissues were screened for CD44 and altered glycosylation (Tn and STn antigens) by different immunoassays (immunohistochemistry, western blot; PLA, double staining immunofluorescence with lectins and antibodies). All procedures were performed under the approval of the hospital’s ethics committee after obtaining informed patient’s consent, being the clinicopathological information obtained from patient’s clinical records.

**TCGA Dataset**

Updated clinical information for 413 TCGA Bladder Cancer (BLCA) cases corresponding to muscle invasive lesions of different stages (T2, T3, T4) and histopathological natures (papillary and non-papillary), including overall survival and disease-free survival information, were obtained from cBioPortal database (https://www.cbioportal.org/). The level-3 RNA-Seq data from 408 TCGA-BLCA cases was retrieved from Broad GDAC FIREHOSE (<http://gdac.broadinstitute.org/>) in March 2021.

**Cell lines and cell culture conditions**

BC cell lines, RT4, 5637 and T24, were acquired from ATCC and cultured in RPMI 1640 GlutaMAX™ medium (Gibco, Thermo Fisher Scientific) supplemented with 10% heat-inactivated FBS (Gibco, Thermo Fisher Scientific) and 1% penicillin–streptomycin (10,000 Units/mL penicillin; 10,000 μg/mL streptomycin; Gibco, Thermo Fisher Scientific). T24 glycoengineered cell models (T24 *C1GALT1*KO and T24 *C1GALT1* KO*/ST6GALNACI*KI) and corresponding controls (T24 *C1GALT1* control carrying a silent mutation and T24 *C1GALT1* control/*ST6GALNAC*1 mock carrying a silent mutation and a mock vector) were generated as described by Peixoto A. *et al.* (1). Transfected cells selection was performed based on puromycin (2 μg/mL, EMD Millipore) resistance. BC cell lines and glycoengineered models were cultured at 37°C in a 5% CO_2_ humidified atmosphere.

**Flow Cytometry**

Cells were detached using Accutase™ Cell Detachment Solution (BD^TM^), fixed with 2% paraformaldehyde (PFA; Sigma-Aldrich), and incubated with rabbit anti-CD44 monoclonal antibody (ab157107, Abcam) using a 1:100 dilution in PBS/2% FBS for 1 h at room temperature (RT). Goat anti-rabbit IgG (H + L) cross-adsorbed secondary antibody Alexa Fluor 488 (Invitrogen) was used for CD44 detection at a 1:300 dilution in PBS/2% FBS for 15 min at RT. Data analysis was performed through CXP Software in a FC500 Beckman Coulter flow cytometer. Results represent the standard deviation of three independent experiments.

**Membrane and Tumor proteins extraction**

Plasma membrane proteins from BC wild type and glycoengineered cell lines were extracted by subcellular fractionation using ultracentrifugation, as previously described (2). Briefly, cultured cells were detached by scrapping with fractionation buffer (20 mM HEPES buffer (pH = 7.4), 10 mM KCl, 2 mM MgCl_2_, 1 mM EDTA and 1 mM EGTA) on ice. The cell suspension was then passed through a 27G needle, left on ice for 20 minutes, and then centrifuged at 720*g* for 5 minutes at 4°C to remove the nuclei. Supernatants were transferred to a new tube and recentrifuged at 10,000*g* for 5 minutes at 4°C to remove mitochondria. Samples were then transferred to polycarbonate centrifuge bottles with cap assemblies and centrifuged for 1 hour at 100,000*g* at 4°C. The pellets were recovered, resuspended in the fractionation buffer, and passed through a 25G needle before a new centrifugation for 45 minutes at 100,000*g* at 4°C. Finally, the plasma membrane-enriched fraction was resuspended in an appropriate volume of TBS with 0.3% SDS. Regarding bladder tumors, total protein was extracted from CD44-STn/Tn expressing areas excised from formalin fixed paraffin embedded tumors using the Qproteome FFPE tissue kit (Qiagen), according to the manufacturer’s instructions.

**Immunoprecipitation**

The Pierce^TM^ Protein G Agarose (Thermo Fisher Scientific) was used to immunoprecipitate CD44 from membrane protein extracts of 5637 and T24 wild type cells and T24 glycoengineered models (100 mg or 500mg of starting material for Western Blotting (WB) and glycoproteomics, respectively) and tumor protein extracts (500mg of starting material). Briefly, agarose beads were blocked with 1% Bovine Serum Albumin (BSA; Sigma-Aldrich) for 1 hour at 4ºC. Prior to immunoprecipitation, membrane protein extracts were cleared with blocked agarose beads for 2 hours at 4ºC, and then, incubated at 4ºC for 2 hours with 3µg (WB) or 6µg (glycoproteomics) of monoclonal anti-CD44 antibody (ab157107; Abcam). The protein-antibody complexes were incubated overnight with newly blocked agarose beads at 4ºC and eluted with SDS Sample Loading Buffer (250 mM Tris–HCl pH 6.8, 8% (w/v), SDS, 0.2% (w/v) bromophenol blue, 40% (v/v) glycerol, 20% (v/v) β-mercaptoethanol). Immunoprecipitated (IP) CD44 was used for further WB and glycoproteomics analysis.

**CD44 separation and proteolytic digestion**

CD44 IP products were resolved by electrophoresis under denaturing conditions using 4-20% gradient precast polyacrylamide gels (Bio-Rad) and bands were excised from gels. Proteins were then reduced with 10 mM 1,4-dithiothreitol (DTT; Sigma-Aldrich) for 45 minutes at 56ºC, alkylated with 10 mM iodoacetamide (Sigma-Aldrich) for 30 minutes in the dark, desialylated with 10U α-neuraminidase [*Clostridium perfringens* neuraminidase Type VI (Sigma-Aldrich)] for 2h at 37ºC and digested overnight at 37ºC with chymotrypsin (25µg/mL; Promega). The proteolytic digests were then analyzed by nanoLC-MS/MS.

**RNAseq for CD44 isoforms identification**

CD44 variants were identified based on a protocol described in detail by Peixoto et al. (3). Summarily, RNA was extracted from 5637 and T24 cell pellets using the RNeasy Plus Mini kit (Qiagen), quantified using Qubit 2.0 Fluorometer (Thermo Fisher Scientific) and RNA integrity was evaluated with Agilent TapeStation (Agilent Technologies). The NEBNext Ultra RNA Library Prep Kit was used to prepare RNA sequencing library for Illumina following manufacturer’s recommendations. After validation and quantification, the sequencing libraries were clustered on one lane of a flow cell, loaded on an Illumina HiSeq 4000 instrument and sequenced using a 2 × 150 Paired End configuration. The HiSeq Control Software was used to analyze the obtained results. The gene hit counts were assessed using the feature Counts from the Subread package v.1.5.2, counting only unique reads that belong to exon regions, and then used for downstream differential expression analysis. A SNP/INDEL and a gene fusion analysis were performed using the Samtools v.1.3.1 program followed by VarScan v.2.3.9 and STAR Fusion v.1.1.0, respectively. For novel CD44 transcripts discovery, Stringtie software was used to extract the transcripts expressed in each cell line. Novel transcripts were pinpointed through the comparison between the reference annotation file and resulting gft file.

**Glycomics**

Bladder cancer cells *O*-glycome was characterized using the Reporter/Amplification method (4), as previously described by us (2, 3). Briefly, cell culture media of semi-confluent cells was supplemented with peracetylated benzyl 2-acetamido-2-deoxy-α-D-galactopyranoside (Sigma-Aldrich, St. Louis, MO, USA) to a final concentration of 150 µM. Following 24 h incubation, glycans were isolated from the conditioned media by filtration using 10 kDa centrifugal filter (Amicon Ultra-4; Merck KGaA, Darmstadt, Germany), followed by solid-phase extraction in Sep-Pak 3 cc C18 cartridges (Waters, Milford, MA, USA). The isolated Bn-*O*-glycosides were then permethylated and analysed by MALDI-TOF-MS on a Bruker UltrafleXtreme mass spectrometer (Bruker Daltonics). Dried samples were resuspended in methanol, mixed (1:1 sample:matrix ratio) with 2,5-dihydroxybenzoic acid (DHB; 10 mg/ml in 50% methanol and 0.1% trifluoroacetic acid; Sigma-Aldrich) and spotted onto a MTP 384 polished steel target plate (Bruker Daltonics). Spectra were acquired in positive ion reflector mode, for a mass range from 540 to 2000 kDa. Then, spectra were subjected to external calibration, using the Peptide Calibration Standard II (Bruker Corporation) combined with α-cyano-4-hydroxycinnamic acid (5 mg/ml in 50% acetonitrile and 0.1% trifluoroacetic acid; Sigma-Aldrich), and internal calibration, using a mass control list constructed by us, considering previous knowledge on bladder cancer *O*-glycosylation.

**Nano-Liquid chromatography-Tandem mass spectrometry**

CD44 IP digests were analyzed by nano liquid chromatography mass spectrometry (nanoLC-MS/MS), exploring an HCD-triggered CID approach. nanoLC-HCD-MS2 was carried out in a Q-Exactive Hybrid Quadrupole-Orbitrap mass spectrometer (Thermo Scientific) coupled to a Ultimate 3000 RSLCnano system (Dionex, Thermo Scientific). Briefly, chymotrypsin digested samples were pre-concentrated in an Acclaim PepMap C18 column (100 Å, 5 mm × 300 µm, i.d. 160454, Thermo Fisher Scientific). Peptide separation was performed in an analytical EASY-Spray column (C18, 100 Å, 2 µm, 75 µm × 500 mm, Thermo Fisher Scientific) with a flow rate of 0.25 µl/min, by mixing the eluent A: 0.1% aqueous formic acid (FA) and eluent B: 0.1% FA in 80% acetonitrile (ACN), with the following gradient: 2 min (2.5% B to 10% B), 50 min (10% B to 35% B), 8 min (35% B to 99% B), and 10 minutes (hold at 99% B). The column was equilibrated with 2.5% B for 17 minutes. The mass spectrometer was operated in the positive ion mode over the *m/z* range 380-1580, and spray voltage was set at 1.9 kV. Full MS settings were the following: 70k resolution (*m/z*=200), AGC target 3*10^6^, maximum injection time 100 ms. The data-dependent parameters were: minimum AGC target 7 × 10^3^, intensity threshold 6.4 × 10^4^, charge state exclusion: unassigned, 1, 8, >8, peptide match preferred, exclude isotopes on, and dynamic exclusion of 20s. The top 10 peaks were selected for HCD fragmentation, using the following MS/MS settings: normalized collision energy (NCE) of 27%, 35k resolution (*m/z*=200), AGC target 2 × 10^5^, maximum injection time 110 ms, isolation window 2.0 *m/z*, isolation offset 0.0 *m/z*, HCD first mass at 110 *m/z*. Mass spectrometer was controlled by Xcalibur 4.0 and Tune 2.9 software (Thermo Scientific). The samples were then run on a nanoLC system coupled to an LTQ-Orbitrap XL mass spectrometer (Thermo Scientific) to allow characterization by collision induced fragmentation (CID). .Liquid chromatography was performed on an EASY-Spray C18 PepMap, 100 Å, 2 µm, 150 mm × 75 µm (Thermo Fisher Scientific) using the same gradient mentioned above. The mass spectrometer was operated in the positive ion mode over the *m/z* range 380-1580. Nanospray voltage was set at 1.9 kV and full scan nominal resolution was 60k (*m/z*=400). CID was triggered from a precursor ion list containing the *m/z* values that presented the HexNAc oxonium ion (*m/z* 204.087 within a ± 0.01 range) in the HCD-MS/MS spectra previously acquired. The 6 most intense ions from the customized parent list were selected for CID fragmentation with NCE = 35%, and MS/MS spectra were acquired in the linear ion trap with an isolation width of 2 Da. Specific parameters were: MS maximum injection time of 500 ms; MS/MS maximum injection time of 50 ms; AGC target 1 × 10^6^ for the Orbitrap and 1 x 10^4^ for LTQ MS^n^ analysis; dynamic exclusion 45s; charge rejection: unassigned and 1. Mass spectrometer was controlled by Xcalibur 3.1 software (Thermo Scientific).

**Bioinformatics for CD44 glycoproteoforms identification**

MS data were first converted to peak lists using Proteome Discoverer version 2.5.0.400 (Thermo Scientific), and then searched against the UniProt Homo sapiens proteome (May 5 2020; 75069 entries), using the MSPepSearch and SequestHT search engines for protein identiﬁcation and the Percolator algorithm v3.05.0 for statistical validation. Prior to the search, the human proteome FASTA database was edited to include CD44 sequences inferred from RNAseq characterization of 5637 and T24 cell lines (**SI Appendix, Table S2**). Searches for HCD tandem spectra were performed with a tolerance of 5 ppm for precursor and 0.02 Da for fragment ions. For CID, a tolerance of 5 ppm was admitted for precursor and 0.6 Da for fragment ions. Chymotrypsin was selected as the proteolytic enzyme and up to two missed cleavages were allowed. A new customized databased composed of high confidence identifications resulting from the initial search was then constructed for definitive CD44 glycoproteoforms identification in glycoproteogenomics settings. For cell lines, carbamidomethylcysteine (+57.0215 Da) was set as a ﬁxed modiﬁcation, while methionine oxidation (+15.9949 Da), protein N-terminal formylation (+27.994 Da), N-terminal acetylation (+42.0106 Da), Asparagine deamidation (+0.9840 Da), ammonia-loss of Cysteine N-terminal (-17.0265 Da) and glutamine to pyro-glutamine modification (-17.0265 Da) were considered as variable modifications. For tumor samples, carbamidomethylcysteine (+57.0215 Da) was elected as a ﬁxed modiﬁcation and, based on previous reports concerning the analysis of FFPE tissues (5), the following oxidative modifications were also included in the variable modifications list: lysine to aminoadipic semialdehyde (-1.0316 Da), arginine to glutamic semialdehyde (-43.0534 Da), proline to pyroglutamic acid (+13.9794 Da), tryptophan to hydroxykynurenin (+19.9898 Da), tryptophan to kynurenine (+3,9949 Da), tryptophan to N-formylkynurenine (+31.9898 Da), threonine to 2-amino-3-ketobutyric Acid (-2.0156 Da), lysine methylation (+14.0156 Da), phenylalanine, Proline, Histidine and Tryptophan hydroxylation (+15.9949 Da) and Phenylalanine, Proline, Histidine and Tryptophan carbonylation (+13.9794 Da) (6). For identification of CD44 glycoproteoforms in cell lines and tumors, the following variable modifications of serine and threonine were also considered: HexNac (+203.0794 Da), NacHexHex 365,1322 Da), HexNac(2) (+406.1588 Da), HexNacNeuAc (494,1748 Da), HexNac(2)Hex (+568.2116 Da), HexNacHexNeuAc (+656,2276 Da), HexNacHexdHex 511,1901 Da), HexNacHexNeuAc(2) (947,3230 Da), HexNac(2) Hex(2)NeuAc (1021.3598 Da). The presence of sialic acids was included to contemplate the possibility of incomplete de-sialylation. Glycoproteomics data was also analyzed using ByonicTM version 2.13.2 (Protein Metrics, Cupertino, CA, USA) using default settings (7). Glycopeptide assignment was confirmed by manual spectra interpretation using the Xcalibur^TM^ software (Thermo Scientific).

**Western and lectin blotting**

Protein extracts and CD44 immunoprecipitates from 5637 and T24 wild type, glycoengineered cells and bladder tumors were separated in 4–20% precast polyacrylamide gels (Bio-Rad) and transferred onto a nitrocellulose membrane (GE Healthcare Life Sciences). STn and Tn expressions were evaluated using the anti-tag-72 antibody [B72.3 + CC49] (1μg/mL, ab199002, Abcam) and biotinylated *Vicia Villosa* lectin (VVA lectin, 1:1000, Vector Laboratories), respectively. CD44 expression was screened with the anti-CD44 antibody (1:5000, ab157107, Abcam). Proteins were blotted with the primary antibody or lectin during 1 hour at RT. The peroxidase affiniPure goat anti-mouse IgG (H+L) polyclonal antibody (1:90,000, ImmunoResearch) was used as a secondary antibody for anti-tag-72 antibody detection, and the goat anti-rabbit IgG (H+L) HPR conjugate antibody (1:60,000; Thermo Fisher Scientific) was used for anti-CD44 antibody detection, both incubated for 30 minutes at RT. The VECTASTAIN® Elite ABC-HRP Reagent (1:10; Vector Laboratories) was used for 15 minutes at RT for analysis of Tn expression. Detection of β-2-microglobulin (B2M) with the recombinant anti-beta 2 microglobulin antibody [EP2978Y] (ab75853, Abcam) followed by incubation with goat anti-rabbit IgG (H+L) HPR conjugate antibody (1:60,000; 30 minutes at RT) was performed as loading control.

**Real-time polymerase chain reaction**

TriPure isolation reagent (Roche Diagnostics GmbH) was used to extract total RNA from BC cells, while RNA from tissues was extracted using the Absolutely RNA FFPE Kit (Agilent), according to the manufacturer’s instructions. RNA reverse transcription and mRNA expression were performed as previously described (8). The relative expression of total CD44 and its isoforms was determined by real-time PCR analysis using TaqMan Gene Expression Assays (total *CD44*: Hs01075864_m1; *CD44v2-v10*: Hs01075866_m1; *CD44v3-v10*: Hs01081480_m1; *CD44v8-v10*: Hs01081475_m1, *CD44s:* Hs01081473_m1; Applied Biosystems) in a 7500 Sequence Detector (Applied Biosystems). *B2M* and Hypoxanthine-guanine phosphoribosyltransferase (*HPRT*) were used for normalization, also as previously described. All samples were run in duplicate and relative mRNA gene expression was calculated with the 2^−ΔCt^ formula.

**Immunohistochemistry**

FFPE bladder tumors and healthy tissue sections were screened by immunohistochemistry for CD44 and Tn and STn antigens, as previously described by Peixoto, A *et al*. (3). Tn antigen expression was evaluated using the biotinylated *Vicia Villosa* (VVA) lectin (Vector Laboratories, 40 mg/mL, 1 hour at 37ºC) and the detection of STn and CD44 antigens were performed using the anti-tag-72 (B72.3 + CC49; Abcam, 0.5 mg/mL, overnight at 4ºC) and anti-CD44 (1:5000, ab157107, Abcam) antibodies, respectively. Lack of cross-reactivity of VVA for blood group A and AB antigens was confirmed using the anti-blood group A monoclonal antibody (HE-193, Thermo Fisher, 1:5, overnight at 4ºC). Sialidase treatment of tissue samples prior to anti-STn probing was also performed to confirm the presence of the glycan (Sigma-Aldrich, 0.2 mg/mL, overnight at 37ºC). CD44 and anti-tag-72 were detected using Novolink Polymer Detection System (Leica) according to manufacturer guidelines. Biotinylated VVA was detected using Streptavidin, Horseradish Peroxidase Conjugate (Thermo Fisher, ready-to-use, 30 minutes, RT) followed by incubation with ImmPACT® DAB Substrate, Peroxidase (Vector, 30:1000, 5 minutes, RT). All images were acquired on a Motic BA310E microscope (Motic) using the Motic Images Plus 3.0 software (Motic).

**Double staining immunofluorescence**

A selection of FFPE tissue sections positive for Tn and CD44 were screened for both antigens through double immunofluorescence to determine colocalization of both epitopes. Briefly, FFPE tissues were deparaffined, hydrated, and exposed to antigen retrieval with EDTA 1mM pH8. Tn antigens were detected using 40 μg/mL FITC-labeled VVA lectin for 2 hours at RT. CD44 antigen detection was achieved using an unlabeled rabbit polyclonal CD44 antibody (Abcam) at 1:250 for 1 hour at RT. An Alexa Fluor 594 anti-rabbit was used for 30 minutes at RT in the dark as a secondary antibody. T24 wild type cells and T24 glycoengineered models were evaluated for T, ST, Tn, STn, and CD44 to detect simultaneous expression between CD44 and these *O*-glycans. Shortly, cells were fixed with 4% PFA for 15 minutes, and then incubated for 1 hour with FITC-labeled VVA lectin (Tn, 0.02 µg/µL), or FITC-labeled PNA lectin (T and ST after desialylation with 70 mU α-neuraminidase, 0.02 µg/µL), or anti-tag-72 (STn, 5 µL/well) and CD44 (1:100). Alexa Fluor 594 anti-rabbit and Alexa Fluor 488 anti-mouse (ThermoFisher Scientific, 1:100) were used to detect CD44 and STn primary antibodies. Nuclear counterstain was performed with 4’,6’-diamidino-2-phenylindole dihydrochloride (2.3x10^-3^ µg/µL; DAPI, Thermo Scientific); for 10 minutes at room temperature in the dark. Fluorescence images were acquired on a Leica DMI6000 FFW microscope using Las X software (Leica).

**Proximity Ligation Assay**

*In situ* proximity ligation assays (PLA) were used for simultaneous detection of CD44 and STn antigens whenever in close spatial proximity at the cell surface in tumors and healthy tissues and glycoengineered bladder cancer cells. Cells were cultured in μ‐Chamber 8-well slides (ibidi), fixed with 4% PFA for 15 minutes. Bladder tumors showing co-localization of CD44 and STn by immunohistochemistry, the healthy urothelium and the other human tissues described in the “Patients sampling and healthy human tissues” sections suggesting CD44 and STn co-localization were evaluated. PFA-fixed cancer cells and tissue sections first undergone antigen retrieval for 15 minutes with boiling citrate buffer pH=6.0 (Vector Laboratories), followed by incubation overnight at 4°C with the conjugated primary antibodies mentioned in the immunohistochemistry section. Ligation and amplification of the PLA signal were achieved using the Duolink PLA Technology kit (Sigma-Aldrich). All slides were incubated with DAPI and mounted using Duolink Mounting Medium. Sialidase treatment prior to antibody probing was used to confirm the specificity of the PLA signals. STn and CD44 negative bladder tumors were used as negative controls. MCR-STn+ glycoengineered cell lines expressing CD44-STn (9, 10) and/or tumor tissues were used as positive controls. The images were acquired on a Leica DMI6000 FFW microscope (Leica Microsystems) using the Las X software (Leica Microsystems).

**siRNA Silencing Assay**

Small interfering RNA (siRNA) reverse transfection was applied to silence *CD44* in 5637, T24, and T24 glycoengineered cells *in vitro*, using a Silencer® Select siRNA targeting CD44 (siRNA ID: s2681, Invitrogen). A Silencer® Select siRNA negative control (4390843, Invitrogen) was also included. Cells were detached and seeded (100,000 cells/well) in a 24-well plate before transfection with lipofectamine RNAiMAX (Invitrogen), according to the manufacturer’s instructions. In brief, Silencer® Select siRNAs and lipofectamine RNAiMAX were diluted in Opti-MEM reduced serum medium (Gibco) and incubated for 5 minutes at RT. Subsequently, cells were incubated with siRNA-lipofectamine complexes for 72 hours at 37°C. Cells were plated in duplicates for each experiment, and the CD44 silencing was confirmed by RT-PCR using the TaqMan Gene Expression Assay Hs01075864_m1.

**Cell proliferation Assay**

The proliferation of 5637, T24 cells and T24 glycoengineered cell models was evaluated in basal and silenced-CD44 conditions, using the cell proliferation ELISA BrdU Kit (Roche Diagnostics GmbH), according to the manufacturer’s instructions. The immunoassay results were monitored at 450 nm using the iMARK™ microplate reader (Bio-Rad). Cell death negative controls composed of 1% Triton-X in a complete cell culture medium were used. The results are presented as the average and standard deviation of three independent assays with three replicates each.

**Invasion Assay**

The invasive capacity of 5637, T24 cells and T24 glycoengineered models, in basal and CD44-silenced conditions, was assessed using Corning BioCoat Matrigel Invasion Chambers (Corning). Briefly, 5×10^4^ cells/mL were plated onto rehydrated invasion inserts according to the manufacturer’s instructions and incubated at 37°C for 24 hours. After removal of non-invasive cells, membranes were washed, and invasive cells were fixed with 4% PFA for 15 minutes. Membranes were then mounted with VECTASHIELD mounting medium with DAPI, and invasive cells were counted in a Leica DM2000 microscope (Leica Microsystems). Three independent experiments were performed, and cells were seeded five times for each experiment. Invasion assays were normalized to cell proliferation average (Invasion Rate).

**Phospho-Kinase Antibody Array**

The relative phosphorylation levels of 37 phosphorylation sites by kinases and 49 receptor tyrosine kinases (RTK) (**SI Appendix, Figs. S7-8**) were determined with the Human Phospho-Kinase/RTK Array kits (ARY003C and ARY001B, respectively, R&D Systems), according to manufacturer’s instructions. Briefly, T24 wild type, glycoengineered cells and corresponding controls were firstly transfected with Silencer Select siRNA targeting CD44 as described above, and then lysed to extract protein from whole cells. For each cell line 300 µg and 600 µg of protein were used for Human Phospho-Kinase and RTK Array kits, respectively. The Amersham ECL Prime Western Blotting Detection Reagent (GE Healthcare Life Sciences) was used as developing reagent. Data analysis was performed through Image Lab Software (Bio-Rad) in a ChemiDoc XRS (Bio-Rad).

**Statistical Analysis**

One-away and Two-way ANOVA followed by Tukey’s multiple comparisons tests and Unpaired T tests were used to determine the different expression patterns of CD44 and its isoforms in BC cells and tissues, as well as in healthy tissues (RT-PCR and Flow cytometry), and to test the effect of CD44 silencing in BC cells and glycoengineered cell models in functional responses (invasion, proliferation). Differences were considered significant for p<0.05. All experiments were performed at least in triplicates and three replicates were conducted for each independent experiment. The results are presented as the average and standard deviation of these independent assays. For the TCGA series, the Shapiro–Wilk normality test was used to determine variable normality. CD44 isoforms log2 (normalized RSEM+1) transformed expression levels were visualized as boxplots. Statistically significant differences between two groups were evaluated using the nonparametric Wilcoxon test, whereas Kruskal-Wallis tests were used for comparison of multiple groups. Spearman correlation analyses were performed to calculate correlation coefficients. Patients were separated into two groups according to the expression levels of each CD44 isoform, using the 25th and 75th percentiles as the cut-off point. Kaplan-Meier (K-M) survival curves were generated to compare the survival between patients with high and low expression levels. The statistical significance between the curves was determined using the log-rank test. The univariate and multivariate Cox proportional hazard regression models were performed to determine independent factors associated with prognosis. All statistical analysis were performed using R software (3.6.2), and *p*-values < 0.05 were considered statistically significant.
